# Haploinsufficiency of eNOS mitigates beneficial effects of maternal exercise on fetal heart development during pregestational diabetes

**DOI:** 10.1101/2025.04.04.647329

**Authors:** Ryleigh Van Neck, Xiangru Lu, Lambertus J. Wisse, Marco C. DeRuiter, Qingping Feng

## Abstract

**Background:** Pregestational diabetes (PGD) increases congenital heart defect (CHD) risk over five-fold. Maternal exercise enhances endothelial nitric oxide synthase (eNOS) activity, benefiting embryos, though its causal role remains unclear. This study investigated eNOS’s role in maternal exercise-mediated protection of fetal heart development in a PGD mouse model.

**Methods:** PGD was induced in eNOS^+/-^ or wild-type (WT) female mice via streptozotocin before breeding with WT or eNOS^+/-^ males. Pregnant females had access to a running wheel for voluntary exercise or remained sedentary. Fetuses were collected at embryonic day (E) 18.5 for genotyping and CHD assessment. E12.5 hearts were analyzed for proliferation, apoptosis, oxidative stress, and eNOS protein levels.

**Results:** Maternal exercise normalized litter size and mortality rates in offspring of diabetic eNOS^+/-^ females but did not reduce CHD incidence in offspring of WT or eNOS^+/-^ females with PGD. CHDs included septal defects, double outlet right ventricle, and valve defects. Exercise increased coronary artery density but not capillary density. Proliferation deficits at E12.5 were restored by exercise, yet oxidative stress remained elevated. Maternal exercise in eNOS^+/-^ dams during PGD did not significantly change eNOS protein levels in both eNOS^+/+^ and eNOS^+/-^ fetal hearts. Offspring genotype did not impact CHD incidence, cell proliferation, apoptosis or oxidative stress.

**Conclusions:** Maternal exercise does not prevent CHDs in PGD offspring of eNOS^+/-^ mice. Its ability to mitigate PGD-induced oxidative stress is eNOS-dependent and essential for improving heart morphology.

**Graphic Abstract:** 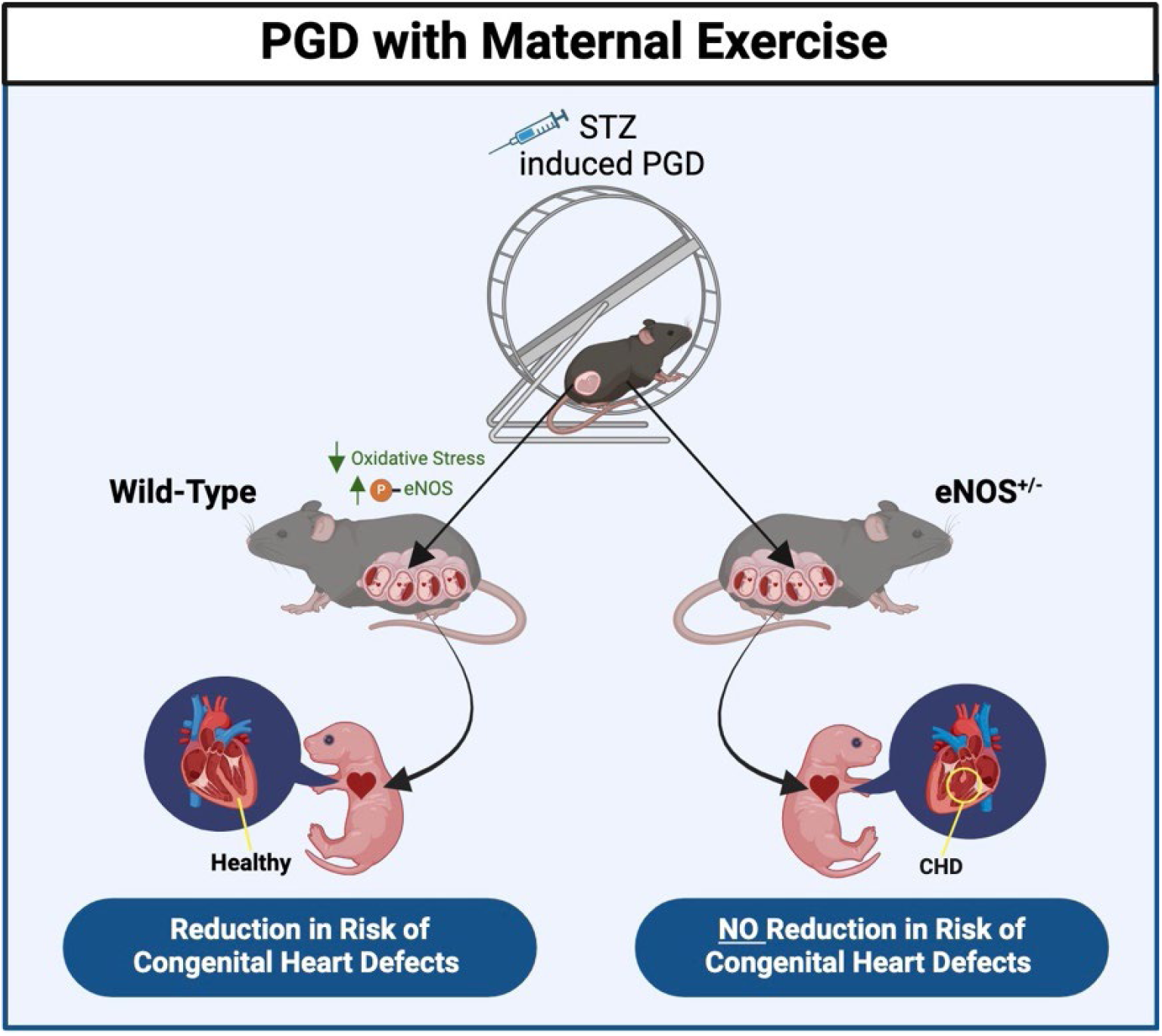

## Background

Congenital heart defects (CHDs) are a leading cause of death in the first year of infant life and arise due to a variety of genetic, maternal and environmental factors (1,2). Pregestational diabetes (PGD) has been identified as an environmental factor that increases the risk of CHDs by more than five times (1–3). The prevalence of CHDs continues to rise globally; although this can be attributed to improved diagnostic screening for asymptomatic conditions, it is also correlated with an increase in diabetes mellitus prevalence in women of reproductive age (4,5). The elevated blood glucose levels associated with PGD interfere with fetal heart development by increasing the production of reactive oxygen species (ROS) and oxidative stress, resulting in abnormalities in cellular apoptosis, proliferation and epithelial-to-mesenchymal transition (EMT), and abnormal placental function, ultimately contributing to alterations in development (6–9).

We have shown that maternal exercise lowers CHD incidence and improves coronary vasculature development in offspring exposed to PGD (10). Specifically, maternal exercise mitigated diabetes-induced changes to myocardial cell proliferation, EMT, and cardiac gene expression (10). Most notably, maternal exercise significantly lowered oxidative stress levels in fetal hearts as demonstrated by a reduction in ROS levels (10). However, the molecular mechanisms responsible for stimulating these beneficial changes remain to be elucidated.

Endothelial nitric oxide synthase (eNOS) is detected in the developing heart as early as E8.5 (11). Nitric oxide (NO) produced by eNOS plays a crucial role in embryonic development, particularly in the formation and maturation of the heart (12–14). NO produced by eNOS contributes to processes essential to heart development including promoting cell survival, cell differentiation, EMT and proliferation. These mechanisms are vital for regulating cardiac septation, valve formation and the development of the coronary vasculature (11,15–18). Furthermore, eNOS-derived NO plays a pivotal role in the fine-tuning of cardiovascular morphogenesis, ensuring the proper structural and functional integration of the fetal heart. Disruption in NO signaling pathways during this critical period can lead to heart defects, highlighting the importance of eNOS in early cardiac development (11,15).

Exercise has been shown to induce eNOS expression and activity in the vascular endothelium in both normal and diabetic subjects (19). Maternal exercise increases eNOS phosphorylation in the fetal heart of offspring exposed to PGD (10). However, the causal relationship between eNOS function and the benefits of maternal exercise on the developing heart in PGD has not been established. In this study, we examined the hypothesis that eNOS haploinsufficiency would prevent the rescuing effects of maternal exercise on heart development in offspring exposed to PGD.

## Methods

### Animals

Mice were used in this study in keeping with the Canadian Council on Animal Care and Use Animal Subcommittee guidelines of Western University, Canada (Protocol #2020-128). Animal suffering and number were minimized in this study. The animals were provided with standard chow and water ad libitum and housed in a 12-hour light/dark cycle. Wild-type (eNOS^+/+^) and eNOS homozygous knockout (eNOS^-/-^, *Nos3^tm1Unc^,* strain #002684) C57BL/6 male and female mice were purchased from Jackson Laboratory (JAX, Bar Harbour, ME, USA). An eNOS^+/-^ line was generated by crossing eNOS^-/-^ females with WT males at 8 weeks old. eNOS^+/-^ offspring were used in all experiments going forward.

Experimental breeding procedures consisted of crossing eNOS^+/-^ females with WT males to produce offspring in the same litter that were WT or eNOS^+/-^. To determine the impact of maternal eNOS^+/-^, an opposing cross was used where WT females were bred with eNOS^+/-^ males to maintain the same litter composition. An eNOS^+/-^ line was used to avoid the spontaneous cardiac defects associated with eNOS^-/-^ lines, such as cardiac septation and coronary artery defects (11,20).

### Induction of Pregestational Diabetes and Voluntary Exercise

WT and eNOS^+/-^ female mice ranging at 8 weeks of age were injected intraperitoneally (IP) with streptozotocin (STZ; Sigma-Aldrich) dissolved in 10 mM sodium citrate for three consecutive days at a dose of 75 mg/kg (10). An additional group of eNOS^+/-^ females without STZ injections served as a control. Blood glucose after 4 hours of fasting was measured one week following the third injection via the tail clip method using the OneTouch Verio2 glucose meter (LifeScan). Mice with blood glucose levels ≥11mmol/L were considered diabetic.

Once diabetes was confirmed, a subset of diabetic WT and eNOS^+/-^ mice were housed in individual cages with access to a voluntary free spinning running wheel (31.92 cm circumference, Columbus Instruments) for a one-week acclimatization period (10). Running wheel revolutions were recorded over 24-hour periods using Multi-Device Channel Interface Software (Columbus Instruments). Daily running distance was calculated by multiplying the number of revolutions by the circumference of the running wheel (Supplemental Table 1). Females that travelled less than 2 km on average per day were excluded from the study. Healthy control eNOS^+/-^ females and diabetic eNOS^+/-^ females were placed in individual cages without a running wheel. Following acclimatization or a one-week-non exercising period, females were mated overnight in cages without a running wheel with males ranging from 8 to 16 weeks of age. The presence of a vaginal plug indicated E0.5. Four-hour fasting blood glucose and body weight were assessed at E0.5 to E18.5 during gestation. There was a total of 8 groups of offspring. Three groups of eNOS^+/-^ dams crossing with WT males gave rise to: control WT and eNOS^+/-^, PGD eNOS^+/+^ and eNOS^+/-^, and PGD+exercise WT and eNOS^+/-^. The 7^th^ and 8^th^ offspring groups were WT and eNOS^+/-^ from WT dams with PGD+exercise crossing with eNOS^+/-^ males.

### Genotyping

Tails and upper limbs were collected to determine fetal genotype. DNA was extracted by incubation with 50 mM sodium hydroxide at 95°C for three hours and subsequently neutralized sing 1 M Tris-HCl. End point polymerase chain reaction (PCR) was performed using DreamTaq Green PCR Master Mix (K1081, Thermo Fisher Scientific, Mississauga, ON). The eNOS^+/-^ genotype was identified using common primer oIMR9358 (5’ CTT GTC CCC TAG GCA CCT CT 3’), eNOS knockout primer oIMR8963 (5’ AAT TCG CCA ATG ACA AGA CG 3’) and WT primer oIMR9357 (5’ <u>AGG GGA ACA AGC CCA GTA GT 3’). PCR products were amplified using a touchdown protocol (Supplemental Table 2 for protocol).</u> The eNOS^+/-^ genotype is indicated by the presence of both a eNOS knockout fragment at 300 bp and the wild-type fragment at 337 bp, where the WT genotype only displays a 337 fragment (Supplemental Figure 1). The PCR products were separated using gel electrophoresis on a 1.7% agarose gel. The proportion of wild-type to eNOS^+/-^ offspring in each maternal condition can be seen in Supplemental Figure 2.

### Cardiac Morphology Assessments

Offspring collected at E18.5 via cesarean section were dissected to isolate the thoracic region of the body and fixed overnight in 4% paraformaldehyde (PFA) at 4°C. The tissue samples were dehydrated using a series of ethanol washes of increasing concentrations (75%; 80%; 95%; 100%), followed by two xylene washes before paraffin embedding. Embedded tissues were cut into 5 µm thick transverse serial sections using a Leica RM2255 microtome (Leica Biosystems) beginning at the level of the aortic arch until the apex. To visualize heart morphology, serial sections were stained with hematoxylin and eosin (H & E; No°2 Hematoxylin Sigma-Aldrich; 1% Eosin in ethyl alcohol VWR) following deparaffinization using xylene (100%; VWR) and serial ethanol rehydration (100% x2, 95%, 75%).

Heart morphology was assessed on a section-by-section basis and the type of CHD was diagnosed by two or more investigators using a light microscope (Zeiss Observer D1).

### Immunohistochemical assessments of Coronary Vasculature

Immunohistochemistry was performed to assess coronary artery and capillary development. Serial sections of E18.5 fetuses were de-waxed and rehydrated using xylene and ethanol. Antigen retrieval was performed in 0.01 M sodium citrate (pH 6.0) for 12 minutes at 94°C in a microwave (BP-111, Microwave Research and & Applications). Anti-α-smooth muscle actin primary antibody 1A4 (A5228, Sigma-Aldrich) was used to identify the smooth muscle component of the coronary arteries at room temperature overnight in a humidity temperature. Biotinylated horse anti-mouse IgG was applied to the tissue for one hour at room temperature. Lectin staining was performed in E18.5 heart sections using biotinylated griffonia simplicifolia lectin-1 (1:50 dilution, Vector Laboratories) to assess capillary density. Colour signals for αSMA and lectin were developed using VECTASTAIN® Elite® ABC-HRP Kit, Peroxidase (Standard) (PK-6100) followed by 3-3’ di-aminobenzidine tetrahydrochloride (DAB, Sigma-Aldrich). Hematoxylin was used as a counterstain.

### Immunofluorescence

Embryonic 12.5 hearts were collected and frozen at −80°C and subsequently embedded in Optimal Cutting Temperature (OCT) Embedding Medium (Tissue-Tek®, Sakura Finetek) at −20°C. Fetal hearts were then sectioned in 8 µm thick sections using the Leica CM 1950 Cryostat (Leica Biosystems).

Cryosections were washed in PBS and fixed in 100% methanol for 10 minutes at −20°C. Cell proliferation, apoptosis and lipid peroxidation were assessed using rabbit anti-phosphorylated histone H3 (pHH3; 1:100; Cell Signaling #9701), rabbit anti-cleaved-caspase-3 (CC3; 1:100; Cell Signaling #9661), and goat anti-4-hydroxynonenal (4-HNE; 1:1000; Applied Biological Materials) respectively.

Primary antibodies were incubated overnight at room temperature followed by goat-anti-rabbit IgG (H+L) Cy-3 conjugated or donkey-anti-goat IgG (H+L) Cy-3 conjugated antibody (1:1500 dilution; Jackson Laboratories) for one hour at room temperature. Slides were then counterstained with DAPI. As a measure of oxidative stress, superoxide levels were quantified using dihydroethidine (DHE; Invitrogen Life Technologies). Sections were incubated in 2 µM of DHE for 30 minutes at 37°C in a light protected humidity chamber and counterstained with nuclear DAPI. All fluorescence signals were imaged using a Zeiss Observer D1 fluorescence microscope and quantification was done using Zen 3.9 Software (Zeiss Germany) or ImageJ (Java 1.8 for PC). The number of proliferating and apoptotic cells was quantified by counting cells positive for pHH3 and CC3, respectively in Zen 3.9 at 20X magnification. The total cell number was quantified using ImageJ to calculate the ratio of proliferating cells to total cells. The rate of apoptotic cells was determined by comparing the number of apoptotic cells to the myocardial area (µm^2^). Lipid peroxidation and superoxide levels were quantified via densitometry at 20X magnification from 3 images per heart and 15 different locations (10,21,22).

### eNOS Protein Levels in E12.5 Hearts

Frozen E12.5 and adult hearts were used for eNOS protein expression assessments via Western blotting. Hearts were homogenized in radioimmunoprecipitation assay (RIPA) buffer (150mM NaCl, 1.0% TritonX-100, 0.5% Na deoxycholate, 0.1% SDS, 50mM Tris pH 8.0, 1mM EDTA). Tissue lysates were collected after centrifugation at 10,000g (4 °C) for 10 min. The Lowry assay (Bio-Rad) was used to assess total protein concentrations and load equal amounts of proteins into a 10% polyacrylamide gel electrophoresis (PAGE). Samples were transferred to a PVDF membrane (Millipore). Membranes were incubated with mouse anti-eNOS antibody (1:1000, BD Biosciences) overnight at 4 °C with gentle agitation followed by mouse-anti-α-actinin antibody (Sigma #A7811, 1:2500 dilution) as a loading control. The eNOS signals were visualized using IRDye® 800CW Goat anti-mouse IgG secondary antibody and IRDye® 680RD goat anti-mouse IgD secondary antibody. Fluorescence signals were visualized using the Odyssey CLx and quantified using Image Studio (Version 3.1, LI-COR Biosciences).

### Statistical Analysis

Data were summarized as mean ± standard error of the mean (SEM). Unpaired Student’s T-test was performed when comparing two groups. One-way analysis of variance (ANOVA) followed by Tukey’s multiple comparison test was used to assess statistical significance between 3 or more groups. The Chi-square test was used to assess CHD incidence and Fisher’s exact test was used to assess craniofacial defect incidence. Data were analyzed using GraphPad Prism, version 10.3 software. Statistical significance was indicated by a *P* value less than 0.05.

## Results

### Bodyweight, fasting blood glucose and running distance during gestation were unchanged by PGD and maternal exercise

Prior to STZ injection, dams in all four maternal conditions underwent a 4-hour fast to establish baseline blood glucose values (Supplemental Figure 3A). Dams with a baseline fasting BG value of >9.0 mmol/L were excluded from the study (n=5). eNOS^+/-^ and eNOS^+/+^ dams were found to have similar basal fasting blood glucose levels. One-week post-STZ injection, both eNOS^+/+^ and eNOS^+/-^ mice had significantly elevated fasting blood glucose values compared to control eNOS^+/-^ dams (*P<*0.05; Supplemental Figure 3B).

Fasting blood glucose was measured at E0.5, E11.5 and E18.5 before fetal collection. Diabetic dams, independent of their genotype and exercise status, demonstrated elevated fasting blood glucose levels compared to control throughout gestation (Figure 1A, *P<*0.05). Additionally, fasting blood glucose was significantly elevated as gestation progressed in dams with PGD (*P<*0.05). Maternal exercise did not improve fasting blood glucose in diabetic eNOS^+/+^ and eNOS^+/-^ dams (Figure 1A). Bodyweight measurements were completed on E0.5, E7.5, E11.5 and E18.5 to ensure dam progression through gestation (Figure 1B). PGD and maternal exercise did not significantly alter maternal body weight at any time point throughout gestation.

**Figure 1.**
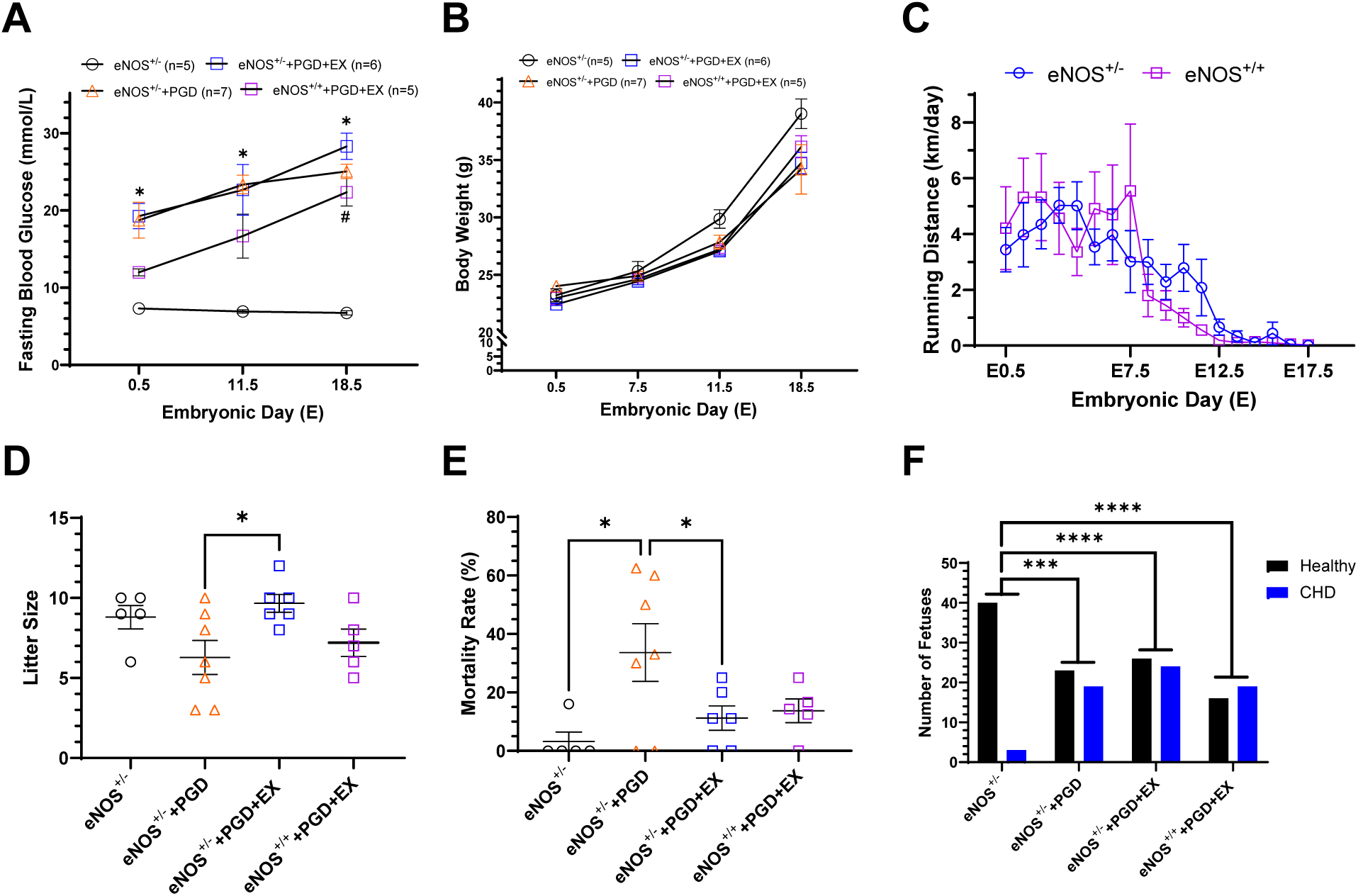
Effects of maternal exercise (EX) on blood glucose, bodyweight, litter size, offspring mortality rate, and congenital heart defects in the offspring of pregestational diabetes (PGD). **A)** Fasting blood glucose levels in eNOS^+/-^ dams with or without PGD and/or EX at E0.5 to 18.5. **B)** Maternal body weight changes during gestation in eNOS^+/-^ dams with or without PGD and/or EX. **C)** Running distance of eNOS^+/+^ and eNOS^+/-^ dams with PDG during gestation. **D)** Litter size of eNOS^+/-^ dams with or without PGD and/or EX at E18.5. **E)** Mortality rate of fetuses of eNOS^+/-^ dams with or without PGD and/or EX at E18.5. **F)** Number of CHD and healthy fetuses of eNOS^+/-^ dams with or without PGD and/or EX at E18.5. The numbers are a combination of both eNOS^+/+^ and eNOS^+/-^ fetuses in each maternal condition. Data represents mean ± SEM and were analyzed using two-way ANOVA followed by Tukey’s test (**A-E**). \**P*<0.05 indicates PGD dams with or without exercise vs eNOS^+/-^ control dams (**A**). #*P<*0.05 indicates eNOS^+/+^+PGD+EX vs eNOS^+/-^+PGD+EX dams (**A**). \**P*<0.05 in **D** and **E**. Fetal CHD numbers were analyzed using Chi-square test with \*\*\**P*<0.001 and \*\*\*\**P*<0.0001 in **F**. eNOS: endothelial nitric oxide synthase, +/-: heterozygous, PGD: pregestational diabetes, EX: exercise.

Between E0.5 and E10.5, diabetic dams travelled approximately 2 to 6 kilometres (km) daily (Figure 1C). Following E10.5, running distance for both eNOS^+/+^ and eNOS^+/-^ genotypes steeply declined due to pregnancy weight gain. The maternal genotype did not alter the running distance throughout gestation (*P*>0.05).

To collect fetuses for assessments of cellular proliferation, apoptosis, oxidative stress and eNOS protein levels at E12.5, a separate cohort of eNOS^+/-^ dams with or without PGD/maternal exercise were studied and their maternal fasting glucose, body weight and running distance were shown in Supplemental Figure 4.

### Voluntary maternal exercise normalized litter size and reduced mortality rate in eNOS^+/-^ dams with PGD

Litter size and mortality rate (% dead) were tabulated at E18.5 during cesarean fetal collection. PGD status impacted litter size in eNOS^+/-^ dams (Figure 1D). Control dams produced litters with a mean size of 8.8±0.7, whereas PGD reduced the mean litter size to 6.3±1.0. Maternal exercise in eNOS^+/-^ dams significantly elevated litter size to a mean of 9.7±0.6 compared to eNOS^+/-^ dams with PGD alone (*P<*0.05). eNOS^+/+^ dams bred with eNOS^+/-^ males did not experience a significant increase in litter size despite maternal exercise (7.2±0.9, *P*>0.05).

Additionally, the fetal mortality rate was significantly elevated by PGD compared to control (Figure 1E, *P*<0.05). Maternal exercise in eNOS^+/-^ dams with PGD significantly reduced the mortality rate (Figure 1E, *P*<0.05). The same trend was observed in fetuses from exercising eNOS^+/+^ dams with PGD, suggesting maternal exercise promotes fetal survival until E18.5 following PGD exposure regardless of maternal genotype.

eNOS^+/-^ fetuses from eNOS^+/-^ dams with PGD experienced a lower mortality incidence than their eNOS^+/+^ counterparts as shown in Table 1 (*P<*0.05). Fetal genotype did not influence mortality risk in fetuses from eNOS^+/-^ control, eNOS^+/-^+PGD+EX, or eNOS^+/+^+PGD+EX dams. These data suggest that eNOS heterozygous insufficiency may be protective against PGD exposure alone.

**Table 1.**
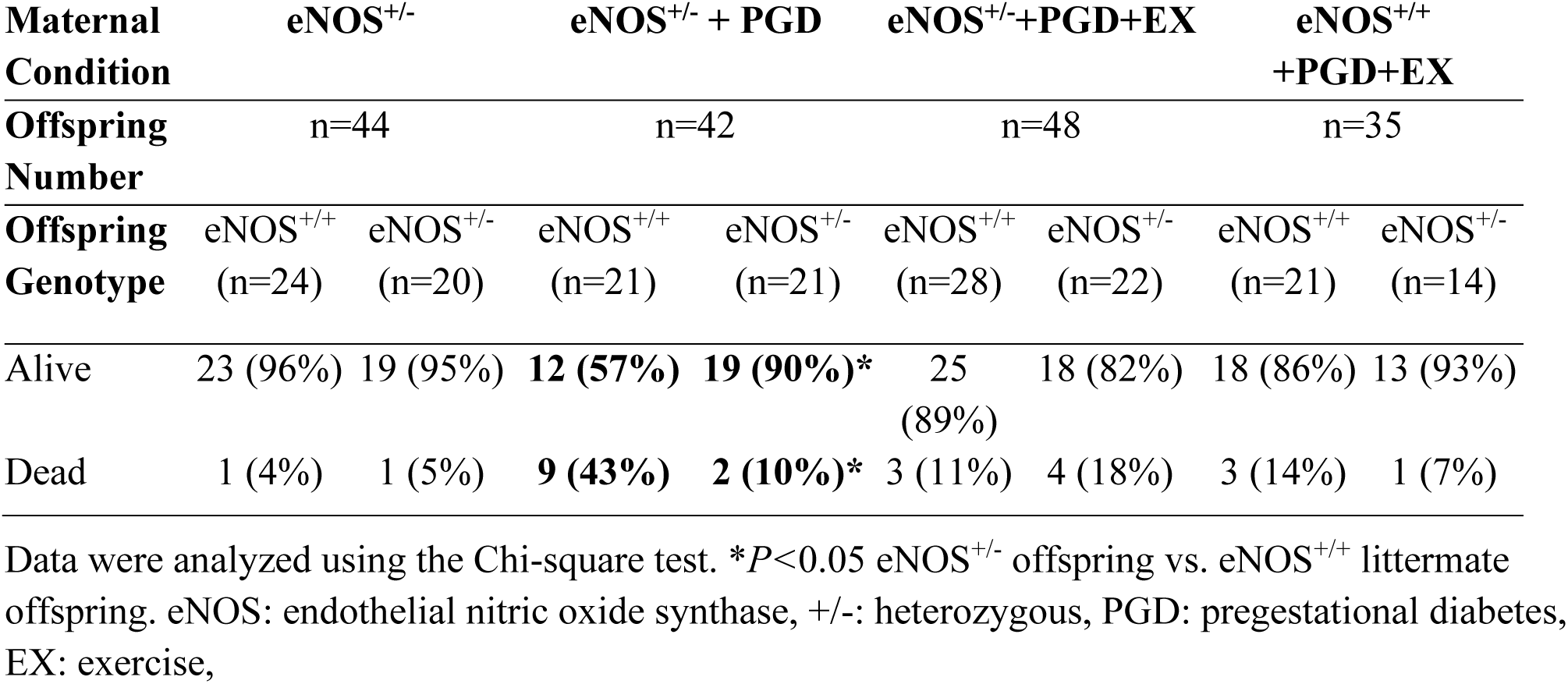
Mortality incidence by genotype in E18.5 fetuses from eNOS^+/+^ or eNOS^+/-^ diabetic dams with or without maternal exercise.

| **Maternal Condition** | **eNOS<sup>+/-</sup>** |  | **eNOS<sup>+/-</sup> + PGD** |  | **eNOS<sup>+/-</sup>+PGD+EX** |  | **eNOS<sup>+/+</sup>+PGD+EX** |  |
| --- | --- | --- | --- | --- | --- | --- | --- | --- |
| **Offspring Number** | n=44 |  | n=42 |  | n=48 |  | n=35 |  |
| **Offspring Genotype** | eNOS<sup>+/+</sup> | eNOS<sup>+/-</sup> | eNOS<sup>+/+</sup> | eNOS<sup>+/-</sup> | eNOS<sup>+/+</sup> | eNOS<sup>+/-</sup> | eNOS<sup>+/+</sup> | eNOS<sup>+/-</sup> |
|  | (n=24) | (n=20) | (n=21) | (n=21) | (n=28) | (n=22) | (n=21) | (n=14) |
| Alive | 23 (96%) | 19 (95%) | **12 (57%)** | **19 (90%)\*** | 25 (89%) | 18 (82%) | 18 (86%) | 13 (93%) |
| Dead | 1 (4%) | 1 (5%) | **9 (43%)** | **2 (10%)\*** | 3 (11%) | 4 (18%) | 3 (14%) | 1 (7%) |

Data were analyzed using the Chi-square test. \* $P < 0.05$ eNOS<sup>+/-</sup> offspring vs. eNOS<sup>+/+</sup> littermate offspring. eNOS: endothelial nitric oxide synthase, +/-: heterozygous, PGD: pregestational diabetes, EX: exercise,

### Voluntary maternal exercise does not reduce CHD incidence in fetuses of eNOS^+/+^ or eNOS^+/-^ dams with PGD

The incidence of CHDs in E18.5 fetuses exposed to PGD with or without maternal exercise was significantly elevated compared to control (Figure 1F, *P<*0.05). Additionally, CHD incidence was not impacted by maternal genotypes as fetuses exposed to PGD, maternal exercise, and differing maternal genotypes (eNOS^+/+^ or eNOS^+/-^) had similar CHD incidences. Therefore, independently of maternal genotype, fetal eNOS haploinsufficiency prevented the rescuing effects of maternal exercise on heart development, which we previously demonstrated (10).

The most diagnosed defects in eNOS^+/+^ and eNOS^+/-^ fetuses exposed to PGD alone (Table 2) were ASD (43% and 48%, respectively), VSD (14% and 14%), ventricular hypertrophy (14% and 19%) and thickening of both the pulmonary (20% and 14%) and aortic valves (10% and 14%) (Figure 2E, F, H, O, P). Other defects observed in this population at lower incidence were AVSD (Figure 2G), DORV (Figure 2M), bicuspid aortic valve (Figure 2N), and cardiac rupture/hemopericardium (Figure 3A-E).

**Figure 2.**
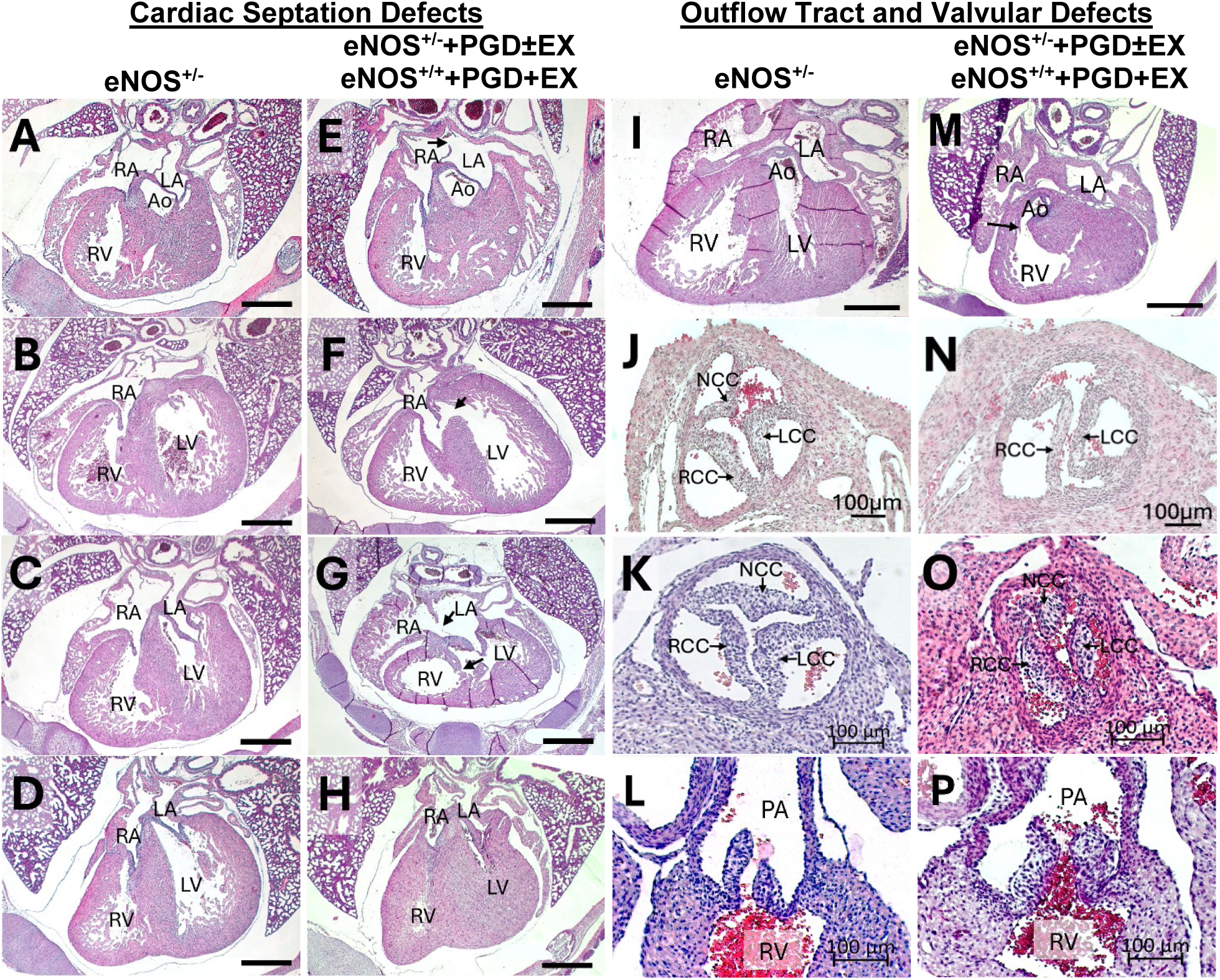
Representative images of cardiac septation defects, myocardial hypertrophy, outflow tract and valve defects in E18.5 eNOS^+/+^ and eNOS^+/-^ offspring exposed to PGD and maternal exercise. **A-C)** Normal cardiac septation morphology from eNOS^+/-^ control dams **D)** Normal myocardial wall thickness, **E)** ASD type 1, **F)** Perimembranous VSD, **G)** AVSD with one common valve, **H)** Myocardial hypertrophy, **I)** Normal right ventricle and aorta morphology, **J-K)** Normal aorta morphology (tricuspid), **L)** Normal aorta connected to LV, **M)** Aorta connected to RV as part of DORV (position more than 50% above RV), **N)** BAV, **O)** thickened aortic leaflets, **P)** thickened pulmonary valve leaflets. eNOS: endothelial nitric oxide synthase, +/-: heterozygous, PGD: pregestational diabetes, EX: exercise, RV: right ventricle, LV: left ventricle, RA: right atrium, LA: left atrium, Ao: aorta, PA: pulmonary artery, NCC: non-coronary cusp, RCC: right coronary cusp, LCC: left coronary cusp. Scale bar is 500 µm in heart images and 100 µm in valve images.

**Figure 3.**
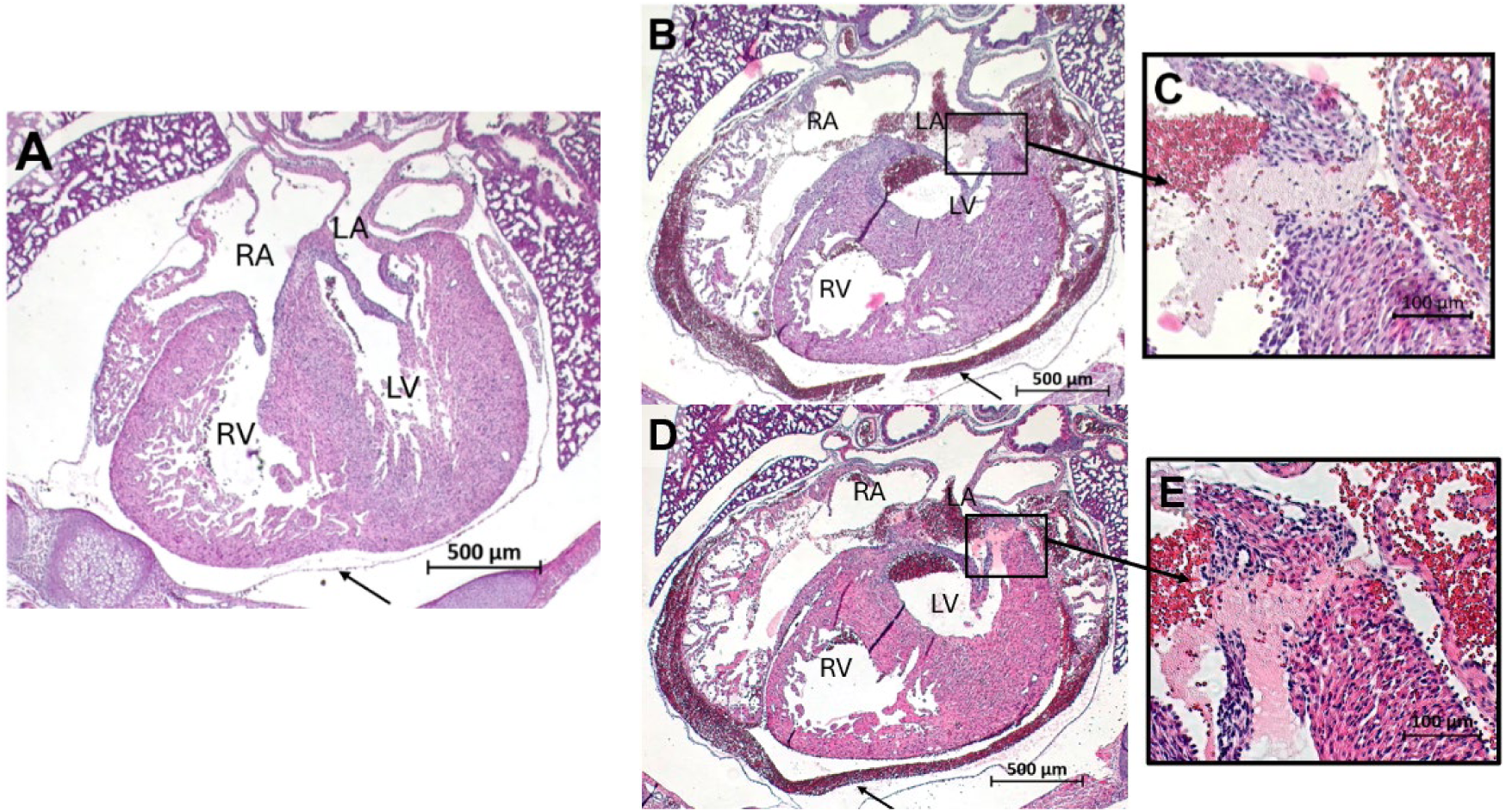
Cardiac rupture and hemopericardium CHD observed in an E18.5 eNOS^+/-^ offspring of an eNOS^+/-^ diabetic dam and maternal exercise. **A)** Normal pericardium morphology represented by the arrow. **B and D)** Cardiac rupture at atrioventricular junction and blood-filled pericardium (hemopericardium) represented by arrow. Scale bar represents 500 µm **C and E)** are enlarged area of **B** and **D**, respectively showing rupture sites and thrombus formation. Scale bar represents 100 µm. **D** and **E** are 25 µm lower than images in **B** and **C**.

**Table 2.** Types of CHDs in fetuses of eNOS+/+ or eNOS+/- dams with PGD, with or without maternal exercise.

| **Maternal Condition** | **eNOS<sup>+/-</sup> (5 litters)** |  | **eNOS<sup>+/-</sup> + PGD (7 litters)** |  | **eNOS<sup>+/-</sup>+PGD+EX (6 litters)** |  | **eNOS<sup>+/+</sup> +PGD+EX (5 litters)** |  |
| --- | --- | --- | --- | --- | --- | --- | --- | --- |
| **Offspring Genotype** | **eNOS<sup>+/+</sup> (n=24)** | **eNOS<sup>+/-</sup> (n=20)** | **eNOS<sup>+/+</sup> (n=21)** | **eNOS<sup>+/-</sup> (n=21)** | **eNOS<sup>+/+</sup> (n=28)** | **eNOS<sup>+/-</sup> (n=22)** | **eNOS<sup>+/+</sup> (n=21)** | **eNOS<sup>+/-</sup> (n=14)** |
| Normal | 22 (92%) | 18 (90%) | 12 (57%)* | 11 (52%)* | 15 (54%)* | 11 (50%)* | 9 (43%)* | 7 (50%)* |
| Abnormal | 1 (4%) | 2 (10%) | 9 (43%)* | 10 (48%)* | 13 (46%)* | 11 (50%)* | 12 (57%)* | 7 (50%)* |
| ASD | 1 (4%) | 2 (10%) | 9 (43%)* | 9 (43%) | 8 (29%)* | 6 (27%) | 10 (48%)* | 6 (43%) |
| VSD | 0 | 0 | 3 (14%) | 3 (14%) | 5 (18%) | 2 (9%) | 2 (10%) | 0 |
| AVSD | 0 | 0 | 0 | 0 | 1 (4%) | 0 | 0 | 0 |
| DORV | 0 | 0 | 1 (5%) | 0 | 2 (7%) | 1 (5%) | 3 (14%) | 1 (7%) |
| Ventricular Hypertrophy | 0 | 0 | 3 (14%) | 4 (19%) | 1 (4%) | 1 (5%) | 6 (29%)*† | 3 (21%) |
| CR/HP | 0 | 0 | 0 | 0 | 0 | 1 (5%) | 0 | 0 |
| BAV | 0 | 2 (10%) | 0 | 3 (14%) | 2 (7%) | 0 | 2 (10%) | 0 |
| Aortic Valve Thickening | 0 | 0 | 2 (10%) | 3 (14%) | 1 (3.5%) | 1 (5%) | 0 | 3 (21%) |
| Pulmonary Valve Thickening | 0 | 0 | 4 (20%) | 3 (14%) | 1 (3.5%) | 1 (5%) | 0 | 3 (21%) |
| Hypoplastic Aorta | 0 | 0 | 0 | 1 (5%) | 0 | 0 | 0 | 0 |

Data were analyzed using the Chi-square test. Comparisons were made between offspring of the same genotype from eNOS<sup>+/-</sup> control dams and experimental groups (\* $P<0.05$ ). Comparisons were made between offspring of the same genotype from eNOS<sup>+/-</sup>+PGD+EX dams vs eNOS<sup>+/+</sup>+PGD+EX dams († $P<0.05$ ). eNOS: endothelial nitric oxide synthase, +/-: heterozygous, PGD: pregestational diabetes, EX: exercise, ASD: atrial septal defect, VSD: ventricular septal defect, AVSD: atrioventricular septal defect, DORV: double outlet right ventricle, CR/HP: cardiac rupture/hemopericardium, BAV: bicuspid aortic valve.

The incidence of these defects was not lowered in eNOS^+/+^ or eNOS^+/-^ fetuses exposed to voluntary maternal exercise, as ASD incidence remained around 50%, regardless of maternal or fetal genotype (Table 2). Notably, the incidence of ventricular hypertrophy was found to be significantly elevated in eNOS^+/+^ fetuses of eNOS^+/+^ dams with PGD and maternal exercise (29%) compared to eNOS^+/+^ fetuses of eNOS^+/-^ dams with PGD and exercise (4%). Ultimately, fetal eNOS heterozygosity did not significantly change the risk of developing any type of CHDs when fetuses were exposed to PGD (Table 2, *P*>0.05).

### Fetal and maternal eNOS haploinsufficiency coupled with PGD and maternal exercise introduced craniofacial defects

No fetuses with craniofacial defects were collected from eNOS^+/-^ control or eNOS^+/+^+PGD+EX dams (Figure 4A). Fetuses exposed to PGD in eNOS^+/-^ mothers with and without maternal exercise had 3 and 5 cases of craniofacial defects, respectively (Figure 4A). An array of craniofacial defects was observed including cleft lip, encephalocele, and exencephaly (Figure 4C & E), however, majority of fetuses had healthy snout morphology (Figure 4B & D). A bilateral cleft lip was observed in our study as the skin failed to close on both sides of the snout (Figure 4C). Encephalocele occurs when the brain and meninges grow through a hole in the fetus’ skull due to a failure of the neural tube to close (Figure 4C) (23). Exencephaly also presents as a protrusion of the brain outside of the skull, however, it is not covered in meninges (Figure 4E). There was no statistical significance in any subgroup comparisons (*P*>0.05) due to low incidences. As PGD with maternal exercise in eNOS^+/+^ dams did not produce any craniofacial defects, the data suggest that maternal eNOS^+/-^ and PGD may contribute to a higher risk of craniofacial defects.

**Figure 4.**
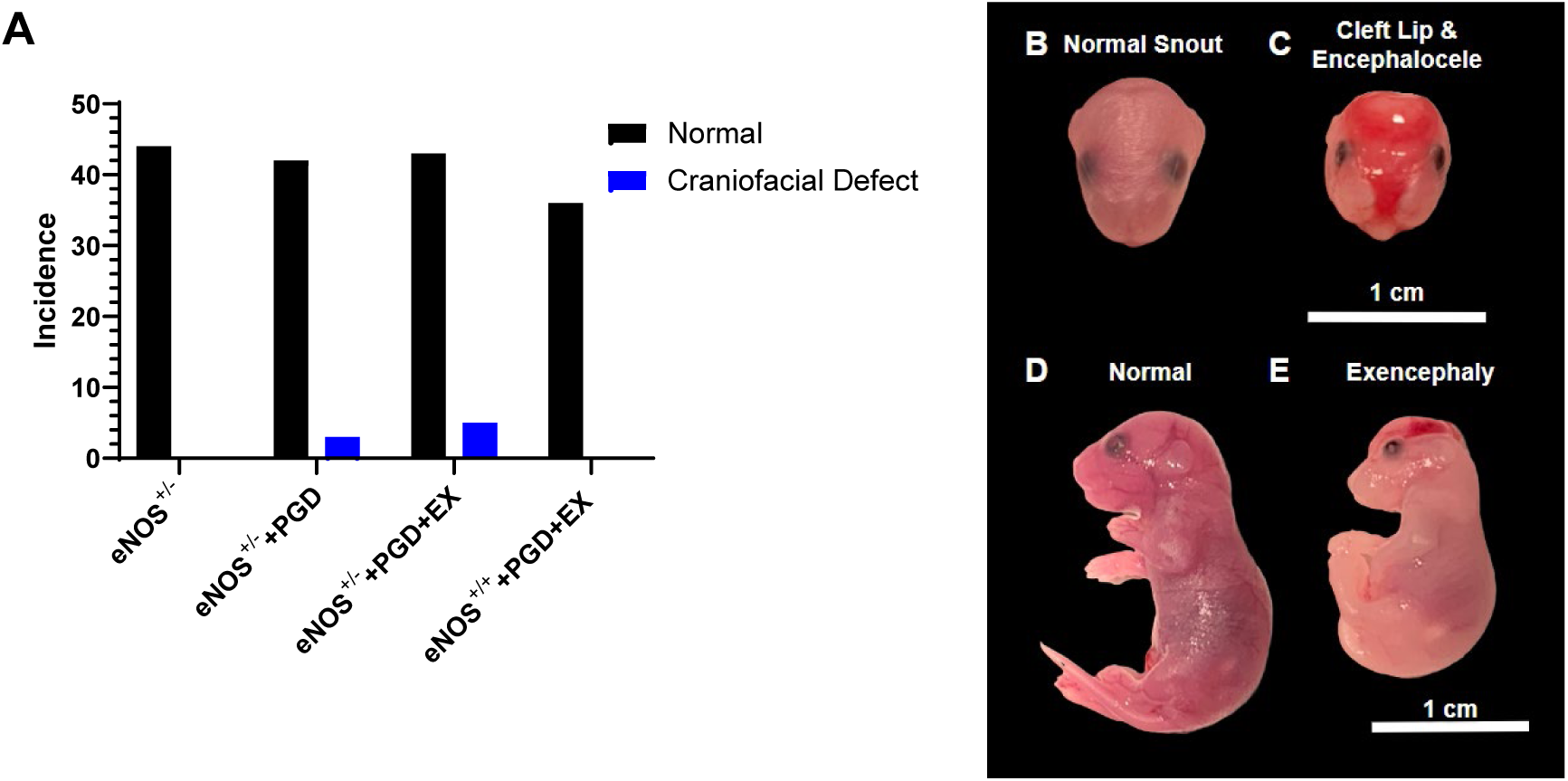
Craniofacial defect incidence and types observed in E18.5 eNOS^+/+^ and eNOS^+/-^ exposed to eNOS^+/-^ dams with PGD with and without maternal exercise. **A)** Incidence of craniofacial defects in offspring exposed to eNOS^+/-^ control dams (n= 44), eNOS^+/-^ +PGD (n=42), eNOS^+/-^+PGD+EX (n=48), and eNOS^+/+^+PGD+EX (n=36). eNOS^+/-^ dams with PGD and maternal exercise produced offspring with the highest risk of craniofacial defect development. **B)** Normal facial and snout morphology. **C)** Cleft lip: failure of the skin to close over the mouth, Encephalocele: brain matter begins to grow outside the skull. **D)** The normal brain is encapsulated by meninges, skull and skin. **E)** Exencephaly: the brain is located entirely outside of the skull. There was no statistical significance by Fishers exact test (*P*=0.0714). eNOS: endothelial nitric oxide synthase, +/-: heterozygous, PGD: pregestational diabetes, EX: exercise.

Fetal genotype alone did not increase the risk of developing a craniofacial defect as evidenced by eNOS^+/+^ and eNOS^+/-^ from control dams displaying no incidences of craniofacial defects (Table 3). Fetuses exposed to PGD alone in eNOS^+/-^ dams displayed no trend in craniofacial defect development correlated with fetal genotype. Additionally, PGD with maternal exercise in eNOS^+/+^ dams displayed no fetal genotype dichotomy in the incidence of craniofacial defects (Table 3).

**Table 3.** Craniofacial defect type and incidence in E18.5 eNOS^+/+^ and eNOS^+/-^ fetuses exposed to diabetic dams without or with maternal exercise.

| **Maternal Condition** | **eNOS<sup>+/-</sup> (5 litters)** |  | **eNOS<sup>+/-</sup> + PGD (7 litters)** |  | **eNOS<sup>+/-</sup>+PGD+EX (6 litters)** |  | **eNOS<sup>+/+</sup> +PGD+EX (5 litters)** |  |
| --- | --- | --- | --- | --- | --- | --- | --- | --- |
| **Offspring Genotype** | eNOS<sup>+/+</sup> (n=24) | eNOS<sup>+/-</sup> (n=20) | eNOS<sup>+/+</sup> (n=21) | eNOS<sup>+/-</sup> (n=21) | eNOS<sup>+/+</sup> (n=28) | eNOS<sup>+/-</sup> (n=22) | eNOS<sup>+/+</sup> (n=21) | eNOS<sup>+/-</sup> (n=14) |
| Normal | 24<br>(100%) | 20<br>(100%) | 20<br>(95%) | 19<br>(90%) | 27<br>(96%) | 18<br>(82%) | 21<br>(100%) | 14<br>(100%) |
| Abnormal | 0 | 0 | 1<br>(5%) | 2<br>(10%) | 1<br>(4%) | 4<br>(18%) | 0 | 0 |
| Exencephaly | 0 | 0 | 1<br>(5%) | 0 | 1 (4%) | 4<br>(18%) | 0 | 0 |
| Encephalocele | 0 | 0 | 0 | 1<br>(5%) | 0 | 0 | 0 | 0 |
| Cleft Lip | 0 | 0 | 0 | 1<br>(5%) | 0 | 0 | 0 | 0 |

Data were analyzed using Fisher's exact test $P=0.0714$ . eNOS: endothelial nitric oxide synthase, +/-: heterozygous, PGD: pregestational diabetes, EX – exercise

### Maternal exercise increased coronary artery abundance in eNOS^+/+^ and eNOS^+/-^ fetuses of eNOS^+/-^ dams with PGD

The smooth muscle cells of the coronary arteries were labelled with α-smooth muscle actin (αSMA) in E18.5 hearts to assess for coronary malformations (Figure 5A-D). PGD did not significantly impact coronary artery abundance, as eNOS^+/+^ and eNOS^+/-^ fetuses exposed to eNOS^+/-^ dams with PGD displayed similar coronary artery abundance to myocardial area ratios (*P*>0.05, Figure 5E). Maternal exercise in eNOS^+/-^ dams with PGD significantly increased fetal coronary artery abundance to myocardial area ratios compared to eNOS^+/-^ control and eNOS^+/-^ dams with PGD conditions (Figure 5E and 5F, *P*<0.05). However, the same trend was not observed in fetuses from exercising eNOS^+/+^ dams with PGD who displayed coronary artery abnormalities that were reflective of fetuses from eNOS^+/-^ control and eNOS^+/-^ with PGD.

**Figure 5.**
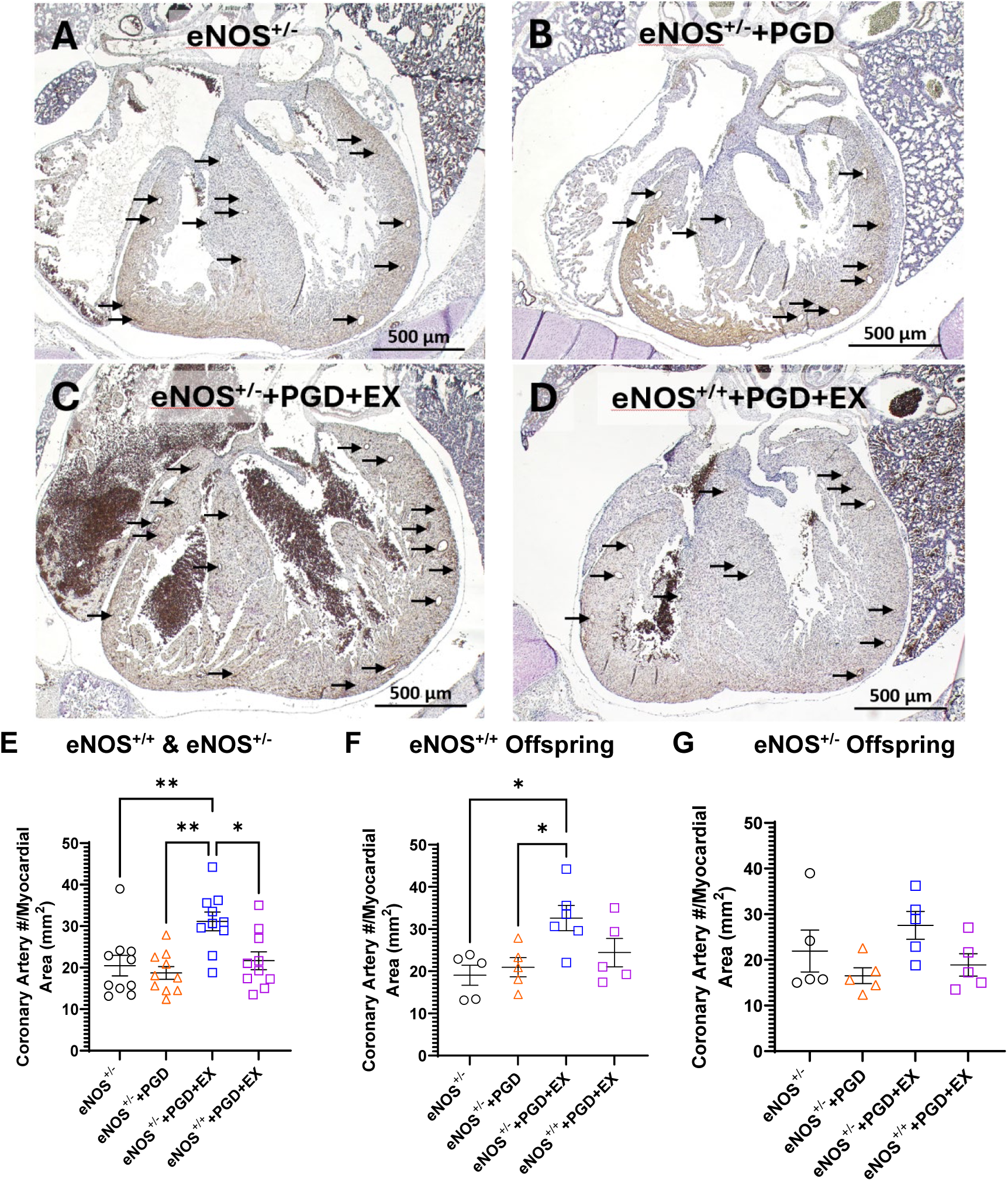
Maternal exercise in eNOS^+/-^ dams with PGD improves coronary artery abundance in E18.5 eNOS^+/+^ and eNOS^+/-^ hearts. **A-D)** Representative images of ventricular coronary arteries in a four-chamber section of E18.5 fetal hearts. Sections are immunostained with α-smooth muscle actin to label coronary arteries. **E)** Average number of coronary arteries counted in three sections for eNOS^+/+^ and eNOS^+/-^ fetuses combined (n=10 hearts per group). **F)** Average coronary artery abundance in eNOS^+/+^ fetuses only (n=5 hearts per group). **G)** The average coronary artery abundance in eNOS^+/-^ fetuses (n=5 hearts per group). Vessels were deemed coronary arteries with a diameter 15-60 µm. Data represent means ± SEM and were analyzed using one-way ANOVA followed by Tukey’s test \**P<*0.05, \*\**P*<0.01. eNOS: endothelial nitric oxide synthase, +/-: heterozygous, PGD: pregestational diabetes, EX: exercise.

The benefits of maternal exercise on coronary artery development were observed in eNOS^+/+^ fetuses (*P*<0.05) but not in eNOS^+/-^ fetuses (Figure 5F and 5G). However, there was no statistical difference in coronary artery abundance to myocardial area between eNOS^+/+^ and eNOS^+/-^ fetuses in any maternal conditions (Supplemental Figure 5, *P*>0.05).

### Capillary density was not impacted by maternal exercise or PGD in eNOS^+/+^ or eNOS^+/-^ fetuses

Capillaries were visualized using biotinylated lectin-1 and capillary density was quantified using three images for the right ventricular myocardium, interventricular septum, and left ventricular myocardium taken at the four-chamber level (Figure 6A-D). Capillary density was normalized to the area of the myocardium and was not significantly altered by changes in maternal genotype, PGD, or maternal exercise (Figure 6E-G). Additionally, there were no significant differences in capillary density relative to the myocardial area between eNOS^+/+^ and eNOS^+/-^ fetuses in each maternal condition (Supplemental Figure 6). However, it should be noted that capillary density in eNOS^+/-^ fetuses tended to be lower than their eNOS^+/+^ counterparts.

**Figure 6.**
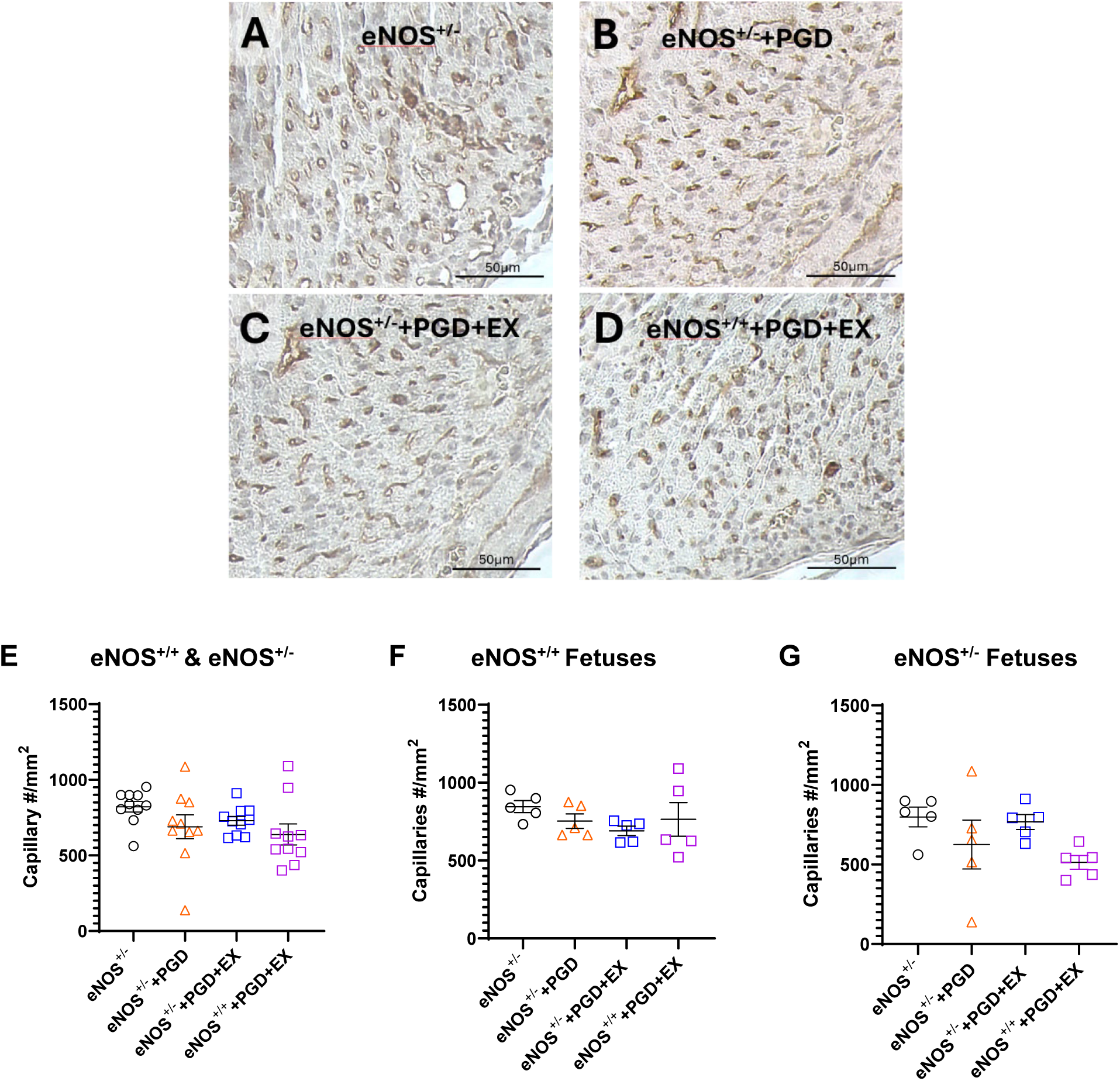
Maternal exercise in eNOS^+/+^ and eNOS^+/-^ dams with PGD does not alter capillary density in E18.5 eNOS^+/+^ and eNOS^+/-^ hearts. **A-D)** Representative images of ventricular capillaries at the four-chamber level of E18.5 fetal hearts. Sections are immunostained with lectin-1 to label endothelial cells. **E)** Average number of capillaries counted in three sections per ventricular structure for eNOS^+/+^ and eNOS^+/-^ fetuses combined (n=10 hearts per group). **F)** Average capillary density in eNOS^+/+^ fetuses only (n=5 hearts per group). **G)** Average capillary density in eNOS^+/-^ fetuses only (n=5 hearts per group). Data represent means ± SEM and were analyzed using one-way ANOVA, *P>*0.05. eNOS: endothelial nitric oxide synthase, +/-: heterozygous, PGD: pregestational diabetes, EX: exercise.

### Maternal exercise normalizes cellular proliferation during PGD in E12.5 eNOS^+/+^ and eNOS^+/-^ hearts

Cellular proliferation was assessed using pHH3 immunofluorescence staining in E12.5 fetal heart cryosections of eNOS^+/-^ control dams (Figure 7A-B). The ratio of proliferating cells to total cells was quantified (Figure 7C). Basal rates of proliferation were not significantly affected by pup genotype (*P*>0.05).

**Figure 7.**
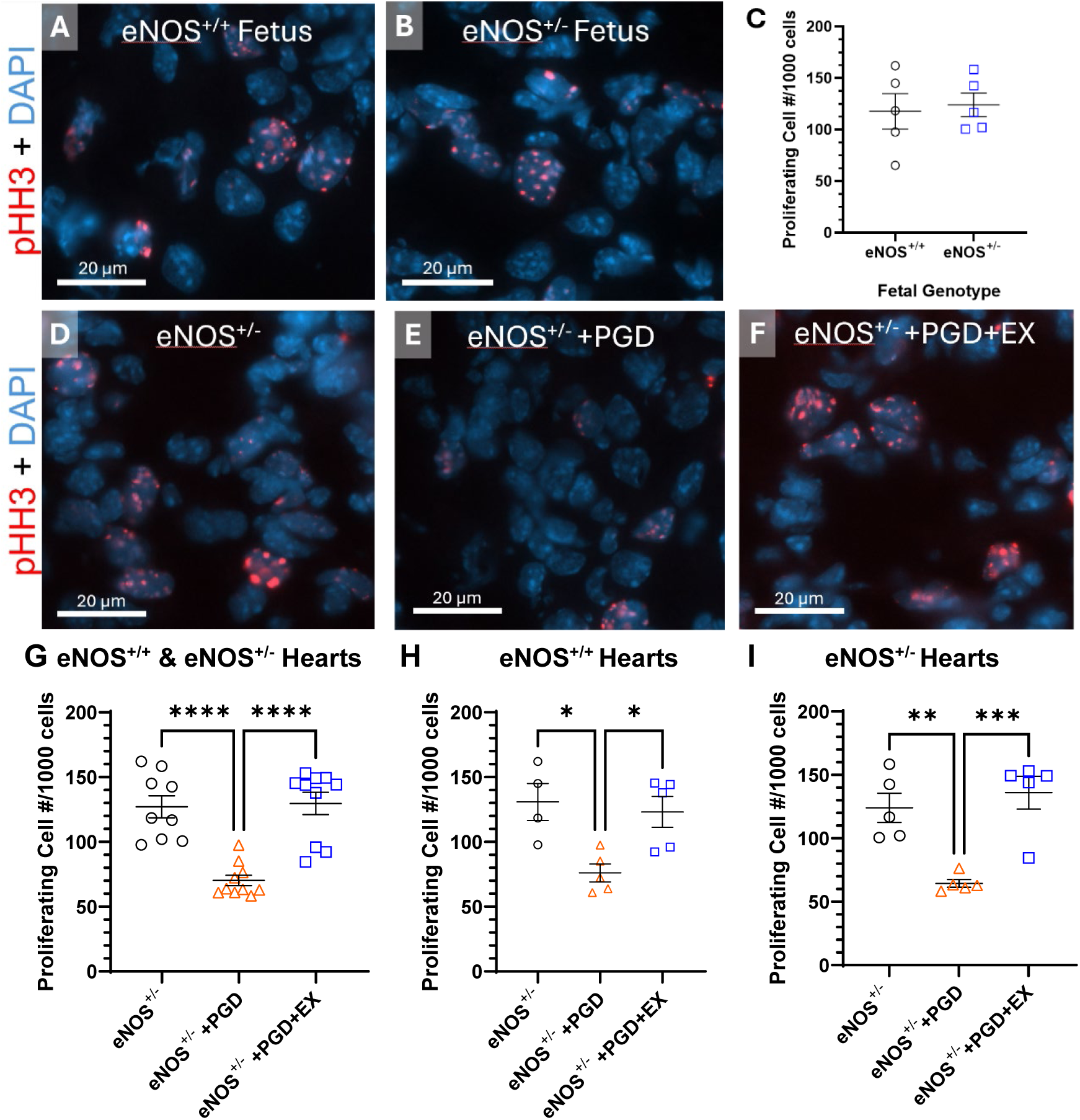
Maternal exercise increased cellular proliferation in E12.5 eNOS^+/+^ and eNOS^+/-^ hearts exposed to PGD. **A-B)** Representative images of pHH3 immunofluorescence in eNOS^+/+^ and eNOS^+/-^ hearts of E12.5 fetuses. **C)** Rates of cellular proliferation per 1000 cells were not significantly different between eNOS^+/+^ and eNOS^+/-^ fetuses. **D-F)** Representative images of pHH3 immunofluorescence in E12.5 fetal hearts from eNOS^+/-^ control, eNOS^+/-^+PGD, and eNOS^+/-^+PGD+EX dams. **G)** Rates of proliferation per 1000 cells were reduced by PGD and rescued by maternal exercise in eNOS^+/+^ and eNOS^+/-^ fetal hearts. The same trend is observed in eNOS^+/+^ hearts **(H)** and eNOS^+/-^ hearts **(I).** Data represent mean ± SEM of n=4-10 hearts per group. \**P<*0.05, \*\**P<*0.01, \*\*\**P*<0.001, \*\*\*\**P*<0.0001 by unpaired Student’s T-test (**C**) or one-way ANOVA followed by Tukey’s test (**G-I**). eNOS: endothelial nitric oxide synthase, +/-: heterozygous, PGD: pregestational diabetes, EX: exercise.

PGD in eNOS^+/-^ dams significantly reduced cellular proliferation in both eNOS^+/+^ and eNOS^+/-^ hearts at E12.5 (Figure 7D-I, *P<*0.05). Maternal exercise in eNOS^+/-^ dams returned cellular proliferation in eNOS^+/+^ and eNOS^+/-^ fetal hearts to control levels (Figure 7D-I, *P<*0.05). Additionally, differences in proliferation between eNOS^+/+^ and eNOS^+/-^ fetal hearts following PGD exposure with or without maternal exercise were assessed. No significant differences in cellular proliferation were found between the two genotypes in either maternal condition (*P*>0.05, Supplemental Figure 7A, B).

### E12.5 eNOS^+/-^ hearts display increased rates of cellular apoptosis compared to eNOS^+/+^ hearts

Cellular apoptosis was assessed using cleaved caspase-3 immunostaining in E12.5 hearts. Representative images are shown in Figure 8A-B. Quantitative analysis showed that eNOS^+/-^ hearts have a greater number of apoptotic cells per mm^2^ than eNOS^+/+^ hearts (Figure 8C, *P*<0.05), suggesting that fetal eNOS heterozygosity increases basal apoptosis in E12.5 hearts.

**Figure 8.**
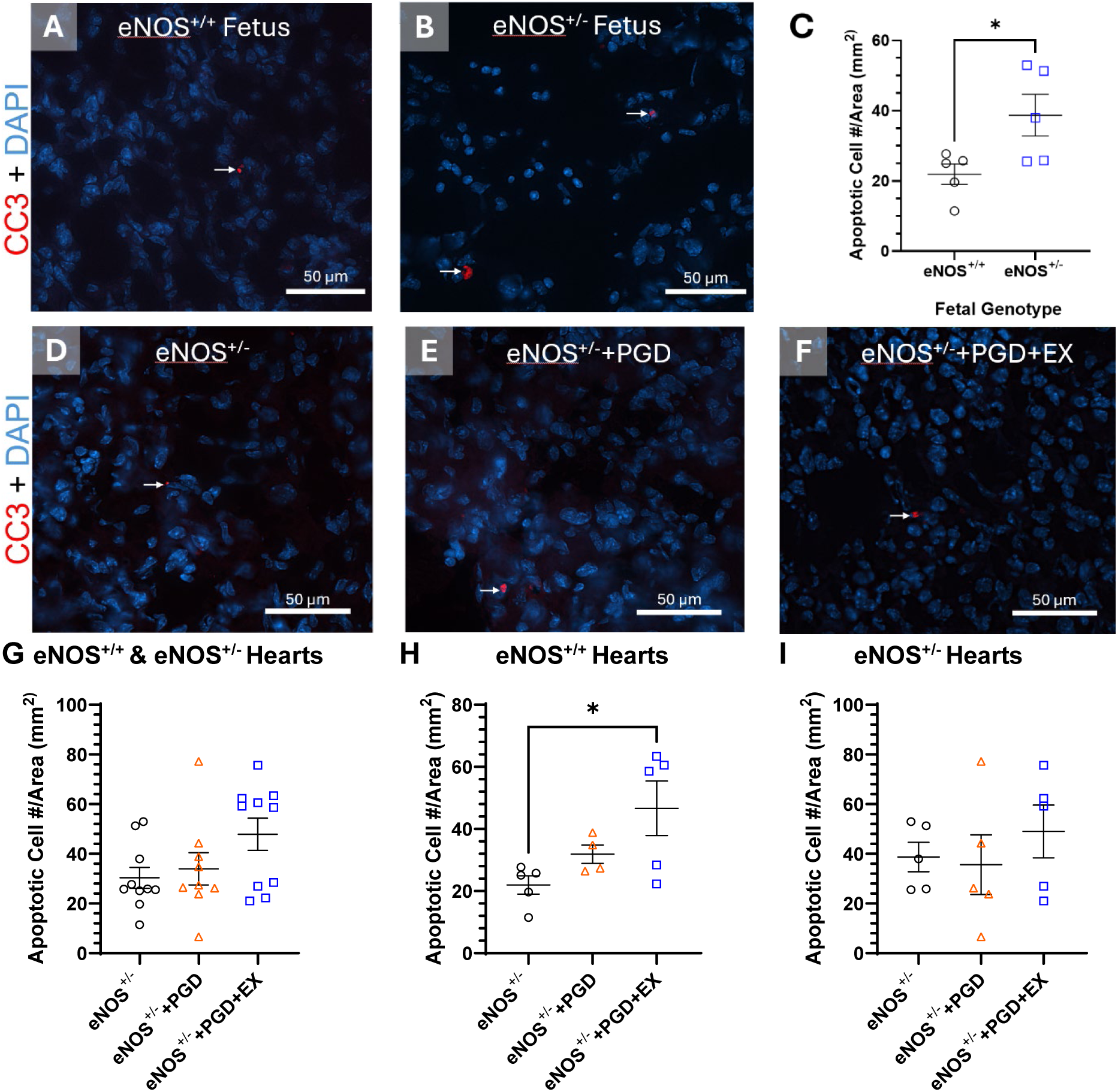
Rates of apoptosis in E12.5 fetal hearts from eNOS^+/-^ dams without and with PGD and/or maternal exercise (EX). **A-B)** Representative images of cleaved caspase-3 immunostaining in E12.5 eNOS^+/+^ and eNOS^+/-^ hearts. **C)** Ratio of apoptotic cells to heart area in fetal eNOS^+/+^ and eNOS^+/-^ hearts of eNOS^+/-^ dams. **D-F)** Representative images of cleaved caspase-3 immunostaining in E12.5 eNOS^+/+^ and eNOS^+/-^ hearts from eNOS^+/-^ control, eNOS^+/-^+PGD, and eNOS^+/-^+PGD+EX dams. **G)** The ratio of apoptotic cells to heart area of eNOS^+/+^ and eNOS^+/-^ fetal hearts. **H)** eNOS^+/+^ fetuses of diabetic dams with exercise showed elevated rates of apoptosis. **I)** The ratio of apoptotic cells to heart area of eNOS^+/-^ fetal hearts. Data represent mean ± SEM of n=4-10 hearts per group and were analyzed using unpaired Student’s T-test (C) or one-way ANOVA followed Tukey’s test. \**P*<0.05. eNOS: endothelial nitric oxide synthase, +/-: heterozygous, PGD: pregestational diabetes, EX: exercise.

### Rates of cellular apoptosis were not significantly altered by PGD with or without maternal exercise in E12.5 eNOS^+/+^ and eNOS^+/-^ hearts

Fetal exposure to PGD did not significantly increase apoptosis rates in E12.5 hearts compared to control (Figure 8D-I). Additionally, maternal exercise in PGD dams did not alter apoptosis in eNOS^+/-^ fetuses or combined eNOS^+/+^ and eNOS^+/-^ fetuses (Figure 8G and 8I) but significantly increased apoptosis in eNOS^+/+^ fetuses (Figure 8H, *P*<0.05).

Differences in apoptosis were investigated between eNOS^+/+^ and eNOS^+/-^ E12.5 hearts exposed to PGD with and without exercise (Supplemental Figure 8). The ratio of apoptotic cells to myocardial area (mm^2^) was not significantly different between the two fetal genotypes (*P*>0.05). eNOS^+/+^ mice exposed to PGD had a mean ratio of 51.0±19.3 apoptotic cells/mm^2^ compared to a mean of 35.6±12.0 apoptotic cells/mm^2^ in eNOS^+/-^ hearts. The same is true in eNOS^+/+^ and eNOS^+/-^ mice exposed to PGD with maternal exercise, as they displayed means of 46.7±8.8 apoptotic cells/mm^2^ and 49.0±10.6 apoptotic cells/mm^2^, respectively. PGD exposure did not significantly alter apoptosis in eNOS^+/-^ fetuses (Figure 8C).

### The impact of fetal eNOS heterozygosity, PGD and maternal exercise on markers of oxidative stress

Two markers of oxidative stress were evaluated: dihydroethidine (DHE) and 4-hydroxynonenal (4-HNE). DHE is a probe for superoxide levels in tissues, whereas 4-HNE is a product of lipid peroxidation visualized using immunostaining. Fluorescence intensity was sampled five times at three separate locations in E12.5 hearts at a constant exposure (150ms for 4-HNE, 90ms for DHE) in each fetal group. Basal superoxide levels were similar between eNOS^+/+^ and eNOS^+/-^ fetal hearts (Figure 9A-C). PGD without and with maternal exercise significantly increased superoxide levels in E12.5 fetal hearts (Figure 10A-D). The same effect was observed in eNOS^+/+^ and eNOS^+/-^ fetuses when analyzed separately (Figure 10E-F). Additionally, no differences in superoxide levels were found between eNOS^+/+^ and eNOS^+/-^ fetuses exposed to PGD or PGD with maternal exercise (*P>*0.05, Supplemental Figure 9).

**Figure 9.**
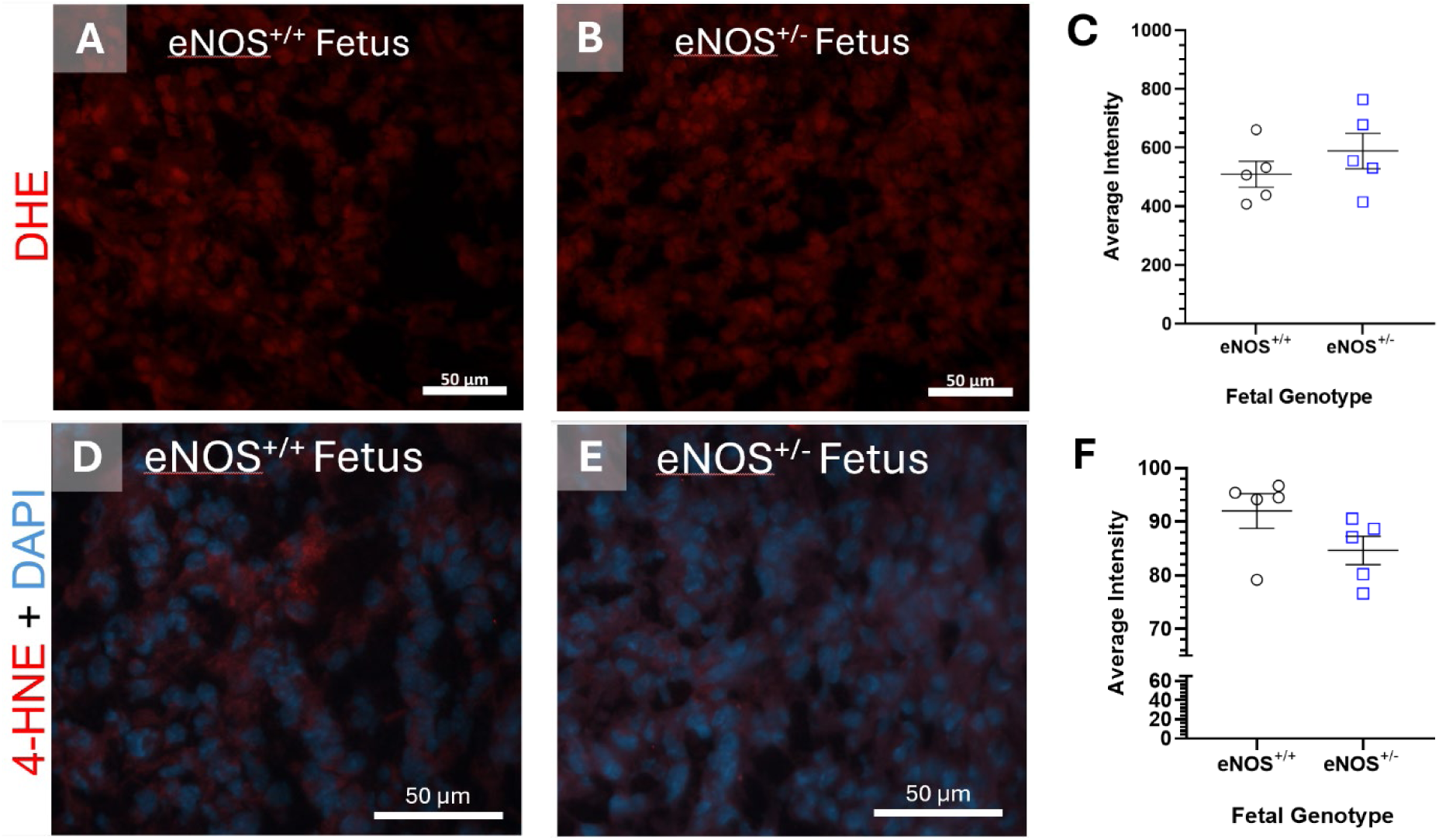
Oxidative stress levels were similar in E12.5 eNOS^+/+^ and eNOS^+/-^ hearts. **A-B)** Representative images of DHE staining in eNOS^+/+^ (**A**) and eNOS^+/-^ (**B**) fetal hearts of eNOS^+/-^ dams. **C)** DHE fluorescence intensity of eNOS^+/+^ and eNOS^+/-^ fetal hearts. **D-E)** Representative images of 4-HNE immunostaining with nuclear DAPI in eNOS^+/+^ (**C**) and eNOS^+/-^ (**D**) fetal hearts of eNOS^+/-^ dams. **F)** 4-HNE fluorescence intensity in eNOS^+/+^ and eNOS^+/-^ fetal hearts. Data represent mean ± SEM of n=5 hearts per group and were analyzed using unpaired Student’s T-test. *P*>0.05. eNOS: endothelial nitric oxide synthase, +/-: heterozygous, PGD: pregestational diabetes, EX: exercise.

**Figure 10.**
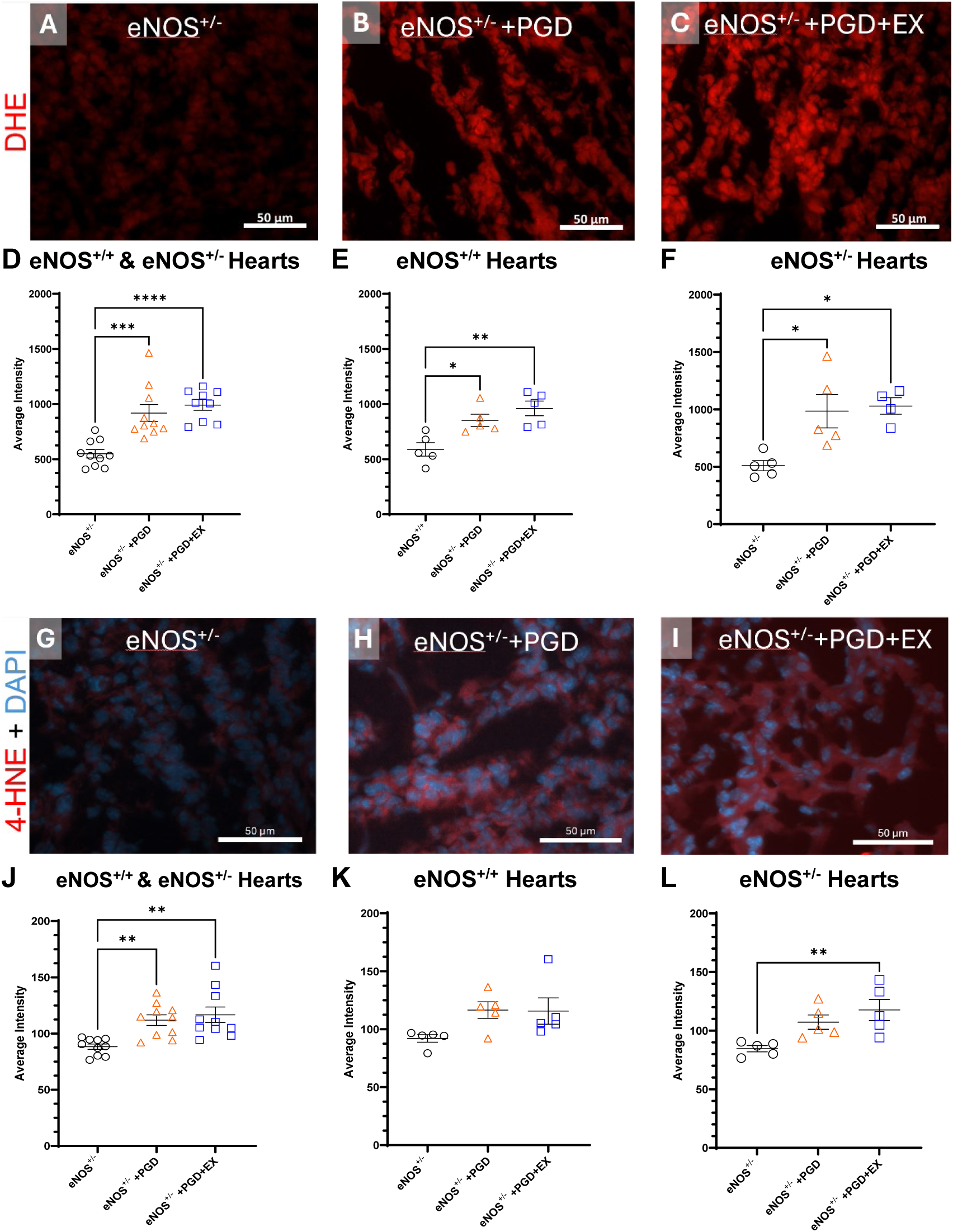
Oxidative stress was elevated by PGD and unchanged by maternal exercise in E12.5 eNOS^+/+^ and eNOS^+/-^ hearts. **A-C)** Representative images of DHE staining in fetal hearts exposed to eNOS^+/-^ dams (**A**), eNOS^+/-^ dams with PGD (**B**), and eNOS^+/-^ dams with PGD and maternal exercise (**C**). **D)** DHE fluorescence intensity in eNOS^+/+^ and eNOS^+/-^ fetal hearts. **E)** DHE fluorescence intensity in eNOS^+/+^ fetal hearts. **F)** DHE fluorescence intensity in eNOS^+/-^ fetal hearts. **G-I)** Representative images of 4-HNE immunostaining and DAPI nuclear stain in fetal hearts exposed to eNOS^+/-^ dams (**A**), eNOS^+/-^ dams with PGD (**B**), and eNOS^+/-^ dams with PGD and maternal exercise (**C**). **J)** 4-HNE fluorescence intensity in eNOS^+/+^ and eNOS^+/-^ fetal hearts. **K)** 4-HNE fluorescence intensity in eNOS^+/+^ fetal hearts. **I)** DHE fluorescence intensity in eNOS^+/-^ fetal hearts. Data represent mean ± SEM of n=5-10 hearts per group and were analyzed using one-way ANOVA followed by Tukey’s test. \**P*<0.05, \*\**P*<0.01, \*\*\**P*<0.001. eNOS: endothelial nitric oxide synthase, +/-: heterozygous, PGD: pregestational diabetes, EX: exercise.

Lipid peroxidation was not significantly different between eNOS^+/+^ and eNOS^+/-^ hearts in the control condition (Figure 9D-F). Exposure to PGD with and without maternal exercise increased oxidative stress in the form of lipid peroxidation in eNOS^+/-^ fetuses but not in eNOS^+/+^ fetuses (Figure 10G-L). Lipid peroxidation levels were not significantly different between eNOS^+/+^ and eNOS^+/-^ hearts from PGD dams with and without exercise (Supplemental Figure 10A, B).

To summarize, eNOS heterozygosity did not increase oxidative stress on its own. PGD without or with maternal exercise increases oxidative stress in the fetal heart. The observed increase in oxidative stress was not dependent on fetal eNOS heterozygosity.

### eNOS protein levels are unaffected by maternal exercise during PGD in E12.5 hearts

The eNOS^+/-^ mouse model was validated by Western blot analysis assessing eNOS protein levels in control eNOS^+/+^, eNOS^+/-^ and eNOS^-/-^ E12.5 hearts (Figure 11A). Protein levels were quantified using densitometry which revealed reduced eNOS protein levels in eNOS^+/-^ hearts compared to eNOS^+/+^ hearts (*P<*0.05), and undetectable expression in eNOS^-/-^ hearts (Figure 11B). Although eNOS^+/-^ animals are deficient in one eNOS allele, protein levels in eNOS^+/-^ hearts were found to be 60% of eNOS^+/+^, rather than the expected 50%.

**Figure 11.**
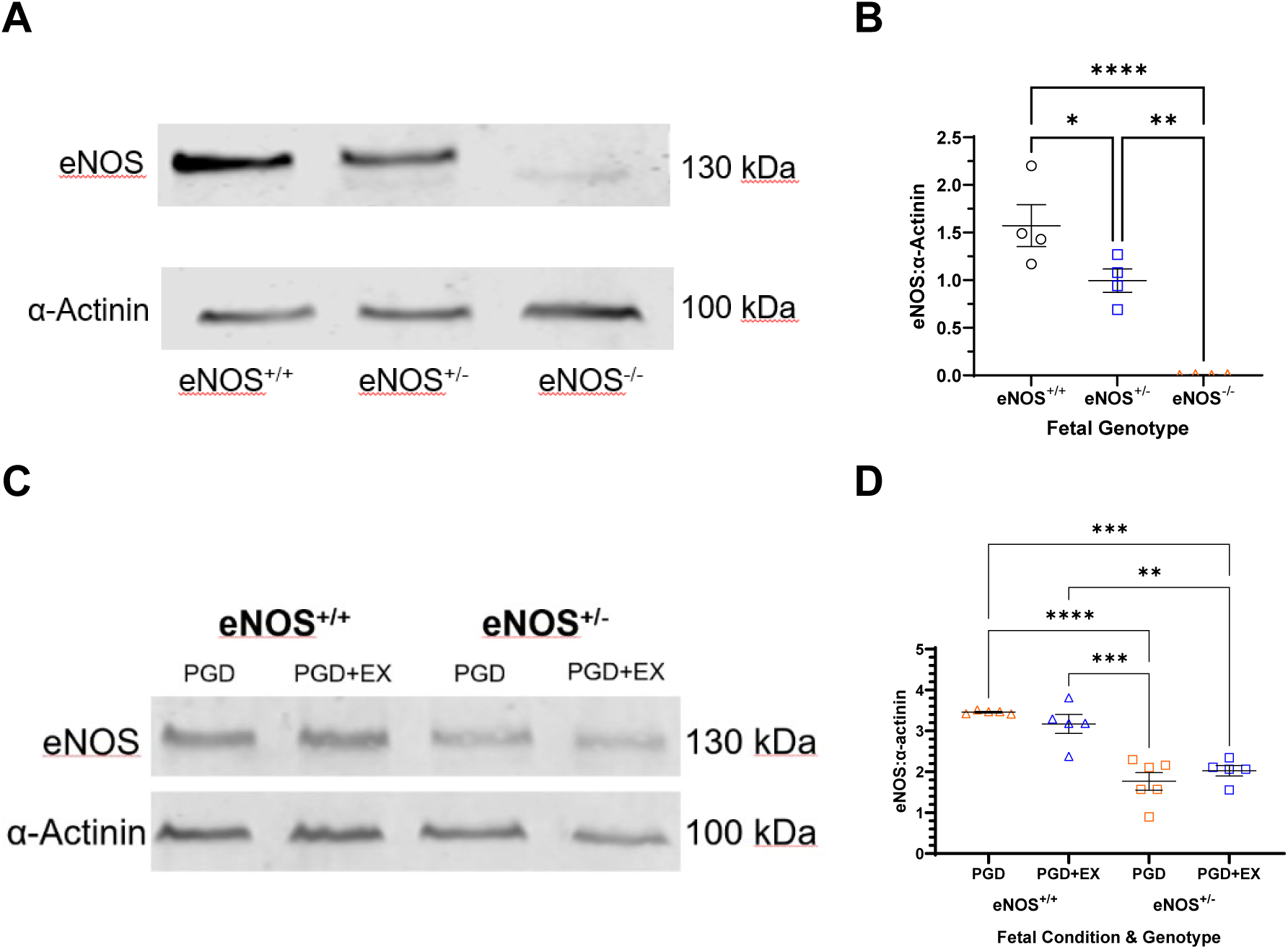
eNOS protein levels in E12.5 fetal hearts of control dams and dams exposed to PGD and maternal exercise. **A)** Representative Western blot depicting eNOS protein levels in eNOS^+/+^, eNOS^+/-^ and eNOS^-/-^ E12.5 hearts of control dams. **B)** Densitometry quantification of the ratio of eNOS to α-actinin. **C)** Representative protein levels from eNOS^+/+^ or eNOS^+/-^ fetal hearts of dams exposed to PGD without and with maternal exercise. **B)** Densitometry quantification of the ratio of eNOS to α-actinin. Proteins depicted are eNOS at 130 kDa and α-actinin at 100 kDa. Data represent mean ± SEM of n=4-6 hearts per group and were analyzed using one-way ANOVA followed by Tukey’s test. *\*P<*0.05, \**\*P<*0.005, \*\*\**P<*0.0005 \*\*\*\**P<*0.0001. eNOS: endothelial nitric oxide synthase, +/-: heterozygous, PGD: pregestational diabetes, EX: exercise.

Similarly, eNOS protein levels in eNOS^+/-^ fetal hearts of eNOS^+/-^ dams exposed to PGD or PGD and maternal exercise were approximately 60% of the eNOS protein in the eNOS^+/+^ hearts (Figure 11C-D). Maternal exercise in eNOS^+/-^ dams during PGD did not significantly change eNOS expression in both eNOS^+/+^ and eNOS^+/-^ fetal hearts (Figure 11C-D, *P>*0.05).

## Discussion

Currently, 16.7 million women of reproductive age (20-49 years) were living with diabetes in 2021 (24). Risk factors for adverse fetal outcomes, such as PGD, can be reduced through effective interventions like glycemic control during pregnancy (3). However, the incidence of CHDs related to maternal diabetes has only slightly decreased by 6% from 1997–2011 compared to earlier years, indicating a need for improved preventative strategies (3). As diabetes prevalence continues to rise, establishing the mechanisms responsible for the benefits of maternal exercise is imperative to improve pre- and antenatal interventions. We aimed to establish a link between eNOS and the benefits of maternal exercise in mitigating CHDs in PGD.

This study demonstrated that maternal exercise did not reduce CHD incidence caused by PGD in an eNOS haploinsufficient environment. However, in wild-type (eNOS^+/+^) mice, we have shown that maternal exercise significantly decreased PGD induced CHD incidence from 59.5% to 25% (10). Our findings show that eNOS haploinsufficient limits the protective effects of maternal exercise during PGD. Interestingly, CHD incidence was similar in fetuses of eNOS^+/+^ and eNOS^+/-^ dams with PGD and exercise, indicating that maternal eNOS may not be the primary mediator of exercise benefits.

Additionally, fetal eNOS heterozygosity did not increase the risk of CHD development during PGD regardless of maternal exercise (Table 2). Hoffman et al. established criteria to classify the severity of CHDs according to the complexity of the intervention required for lifetime management of the defect (25). According to these criteria, ASD, VSD, and valve defects (leaflet thickening, BAV) are less severe than more complex defects like DORV and AVSD. Previous research showed that maternal exercise during PGD reduced the incidence of more severe CHDs, like DORV and AVSD (10). However, the current study found no difference in the incidence of more severe to less severe defects across maternal conditions (Supplemental Table 3). Furthermore, while prior studies have reported additional severe defects like hypoplastic heart syndrome (10), which were not observed in this study. Differences in blood glucose levels between study cohorts throughout gestation may explain these discrepancies. Additionally, the fetal genotype did not influence the risk of developing CHDs, reenforcing the conclusion that fetal eNOS alone does not independently drive the benefits of maternal exercise.

Cardiac and vascular defects arise when the normal steps of cardiogenesis are impeded. E12.5 hearts exposed to PGD showed less pHH3 immunostained cells, indicating fewer cells undergoing proliferation. Maternal exercise restored cellular proliferation impaired by PGD to control levels in our study regardless of fetal genotype. We have previously shown that PGD impairs cellular proliferation in fetal hearts and maternal exercise restores cellular proliferation to control levels (10). The improved cellular proliferation was attributed to increased expression of *Cyclin D1*, a marker of cell proliferation, and cardiogenic transcription factor *Gata4* that is critical for the driving proliferation at key stages of cardiac development, such as septation (10,26,27). In the present study, eNOS haploinsufficiency did not impede the maternal exercise benefits to cellular proliferation. Notably, maternal exercise has been shown to normalize the dysregulated miRNAs enriched in cell proliferation (28). Our data suggest that maternal exercise may rescue cell proliferation in the fetal heart via eNOS independent signaling pathways, such as miRNA expression networks.

Maternal exercise during PGD did not alter eNOS protein levels in E12.5 fetal hearts. This aligns with findings in adult male eNOS^+/-^ mice, where exercise did not change eNOS mRNA or protein levels in blood vessels, though eNOS expression was increased in the left ventricular myocardium (29). In wild-type fetal hearts, maternal exercise during PGD did not significantly affect eNOS protein levels but did increase in eNOS phosphorylation (10). These data suggest that the protective effects of maternal exercise during PGD are not driven by an overall increase in eNOS protein expression. Notably, eNOS phosphorylation at serine 1177 increases its activity. Whether maternal exercise increases eNOS phosphorylation in fetal hearts from eNOS^+/-^ dams during PGD remains be investigated in future studies.

A major contributor to CHD pathogenesis is oxidative stress (9,30,31). Our study found oxidative stress levels were elevated in E12.5 fetal hearts exposed to PGD, consistent with previous research (7,10,32). This increase was evident through higher superoxide and lipid peroxidation levels. However, maternal exercise did not reduce oxidative stress in E12.5 eNOS^+/+^ or eNOS^+/-^ hearts of fetuses from eNOS^+/-^ dams during PGD. Previously, we demonstrated that maternal exercise lowers superoxide and lipid peroxidation levels in eNOS^+/+^ fetal hearts during PGD (10). This discrepancy in our current findings may be due to reduced NO availability caused by eNOS heterozygosity, which compromises its antioxidant properties. NO can efficiently transfer from maternal to fetal circulation via the placenta, a shared pool of NO likely exists regardless of fetal genotype. Exercise has been shown to increases NO production in wild-type adults, but not in eNOS^+/-^ animals (19,29). NO mitigates Fenton-mediated oxidative stress through several mechanisms, including direct scavenging, inhibition of peroxide reactions, and neutralization of superoxide. For example, NO can bind to ferrous iron in softer ligand fields, such as heme complexes, forming a metal-nitrosyl complex that prevents peroxide-iron interactions and ROS production. In addition, NO reacts with superoxide to form peroxynitrite, which is rapidly converted to nitrate, reducing ferric iron (Fe³⁺) and halting ROS formation (33). In the present study, a lower NO level in eNOS^+/-^ dams may have been insufficient to counteract ROS production induced by PGD (6,34).

In adults, exercise increases ROS production, which paradoxically enhances redox capacity by upregulating antioxidant enzymes such as SOD, catalase and GPx in cardiac and skeletal muscles. This adaptive response strengthens antioxidant defenses, protecting against ischemia-reperfusion injury (35,36). Additionally, eNOS-derived NO plays a crucial role in downstream signaling, influencing gene expression (37). Fukai et al. reported that exercise-induced eNOS activation correlates with increased SOD expression in adult vasculature, leading to a subsequent reduction in ROS (38). Recent studies in humans have revealed that a specific isoform of SOD, SOD3, is upregulated by maternal exercise in a T2DM model in the placenta which subsequently leads to activation of the AMPK/TET pathway in the fetal liver (39). Furthermore, Kusuyama et al. demonstrated that maternal exercise induces epigenetic modifications in utero, leading to long term improvements in fetal liver function (39). These findings highlight the need for further research into crosstalk between placenta and fetal organs to better understand how maternal exercise promotes fetal heart development.

## Conclusions

In summary, maternal exercise during PGD in eNOS^+/-^ dams does not alleviate PGD-induced oxidative stress or the incidence of CHDs in fetuses. Our findings suggest that while maternal exercise can influence fetal cardiac oxidative stress in utero, this effect depends on eNOS. Since eNOS haploinsufficiency diminishes the benefits of maternal exercise on fetal heart development, our study highlights a critical role of eNOS in mitigating CHD risk during PGD by maternal exercise.

## Acknowledgements

RVN designed the research, performed the experiments, analyzed data, drafted and revised the manuscript. XL aided in study design, performed the experiments, analyzed data, and revised the manuscript. LW and MD aided in the examination of fetal heart morphology. QF conceived and designed the research, interpreted the results and revised the manuscript. RVN, XL, and QF are guarantors of this work and had full access to all the data in the study and therefore take responsibility for the integrity of the data and the accuracy of its analysis. All authors read and approved the manuscript. We thank our volunteers: Ivana Du, Thomas Pan, Adam Yanagi, and Nathan Lee for their contributions to the generation of histological slides. The graphic abstract was created in BioRender. Van neck, R. (2025) https://BioRender.com/y14q870.

## Non-Standard Abbreviations and Acronyms

ASD: Atrial septal defect
AV: Atrioventricular
AVSD: Atrioventricular septal defect
BAV: Bicuspid aortic valve
CC3: Cleaved caspase-3
CHD: Congenital heart defect
DHE: Dihydroethidium
DM: Diabetes mellitus
DORV: Double outlet right ventricle
EMT: Epithelial to mesenchymal transition
eNOS: Endothelial nitric oxide synthase
GPx: Glutathione peroxidase
4-HNE: 4-hydroxynonenal
LA: Left atrium
LV: Left ventricle
miRNA: Micro-RNA
PGD: Pregestational diabetes
pHH3: Phosphorylated histone H3
ROS: Reactive oxygen species
RA: Right atrium
RV: Right ventricle
SOD: Superoxide dismutase
STZ: Streptozotocin
αSMA: α-smooth muscle actin
T1DM: Type 1 diabetes mellitus
T2DM: Type 2 diabetes mellitus
VSD: Ventricular septal defect
Wnt: Wingless/Integrated
eNOS+/+: Wild type
eNOS+/-: eNOS heterozygous

