## Supplemental tables and figures for "Haploinsufficiency of eNOS mitigates beneficial effects of maternal exercise on fetal heart development during pregestational diabetes"

### Supplemental Materials

**Supplemental Table 1. Running distance for diabetic dams used in E18.5 fetal collection.** Data was collected using the mouse home cage running wheel from Columbus Instruments. Distance was calculated using:

$$Running\ Distance\ (km) = \frac{(31.91cm \times Wheel\ Revolutions)}{100\ 000}.$$

| Mouse ID | Wheel Revolutions in 24 hours | Mean Distance Per Day (km) | Mean Distance During Acclimatization (km) |
|---|---|---|---|
| | **eNOS$^{+/+}$ Females +PGD+EX** | | |
| V88 | 18248 | 5.85 | 7.80 |
| | 5964 | 1.91 | |
| | 17670 | 5.66 | |
| | 26051 | 8.35 | |
| | 33374 | 10.69 | |
| | 34736 | 11.13 | |
| | 34449 | 11.04 | |
| V94 | 9296 | 2.98 | 4.96 |
| | 7865 | 2.52 | |
| | 13540 | 4.34 | |
| | 15357 | 4.92 | |
| | 17449 | 7.46 | |
| | 16753 | 5.37 | |
| | 22339 | 7.16 | |
| V96 | 6022 | 1.93 | 6.98 |
| | 17814 | 5.71 | |
| | 19553 | 6.27 | |
| | 24456 | 7.84 | |
| | 29646 | 9.50 | |
| | 23053 | 7.39 | |
| | 31923 | 10.23 | |
| V101 | 26589 | 8.52 | 8.36 |
| | 15673 | 5.02 | |
| | 14098 | 4.52 | |
| | 22617 | 7.25 | |
| | 32179 | 10.31 | |
| | 35769 | 11.46 | |
| | 41256 | 13.22 | |
| V111 | 23727 | 7.60 | 10.03 |
| | 16262 | 5.21 | |
| | 24205 | 7.76 | |
| | 26580 | 8.52 | |
| | 44051 | 14.12 | |
| | 38696 | 12.40 | |
| | 45502 | 14.58 | |

**Supplemental Figure 1 Continued…**

| Mouse ID | Wheel Revolutions in 24 hours | Mean Distance Per Day (km) | Mean Distance During Acclimatization (km) |
|---|---|---|---|
| | **eNOS$^{+/-}$ Females +PGD+EX** | | |
| V34 | 24300 | 7.79 | 9.96 |
| | 18887 | 6.05 | |
| | 23369 | 7.49 | |
| | 35709 | 11.44 | |
| | 35709 | 11.44 | |
| | 35709 | 11.44 | |
| | 53147 | 17.03 | |
| V35 | 14998 | 4.81 | 6.96 |
| | 19684 | 6.58 | |
| | 24702 | 7.92 | |
| | 24702 | 7.92 | |
| | 24702 | 7.92 | |
| | 24702 | 7.92 | |
| | 26682 | 8.55 | |
| V36 | 15699 | 5.03 | 4.61 |
| | 10747 | 3.44 | |
| | 14085 | 4.51 | |
| | 17217 | 5.52 | |
| | 17217 | 5.52 | |
| | 17217 | 5.52 | |
| | 14151 | 4.53 | |
| V39 | 9471 | 3.03 | 3.47 |
| | 9471 | 3.03 | |
| | 5356 | 1.72 | |
| | 8490 | 2.72 | |
| | 15348 | 4.92 | |
| | 15348 | 4.92 | |
| | 15476 | 4.96 | |
| V44 | 17102 | 5.48 | 4.91 |
| | 11993 | 3.84 | |
| | 13238 | 4.24 | |
| | 17401 | 5.58 | |
| | 11993 | 3.84 | |
| | 11993 | 3.84 | |
| | 16861 | 5.40 | |
| V45 | 15472 | 4.96 | 4.81 |
| | 10202 | 3.27 | |
| | 10202 | 3.27 | |
| | 10202 | 3.27 | |
| | 11879 | 3.81 | |
| | 15027 | 4.82 | |
| | 22503 | 7.21 | |

**Supplemental Table 2. End-point PCR cycling parameters used for genotyping procedures used to identify eNOS$^{+/-}$ fetuses.**

PCR was run using the MyCycler™ Thermal Cycler (Bio-Rad Laboratories) based on Protocol 23416 (Jackson Laboratories).

| Step | Temperature (°C) | Time | Number of Cycles |
|---|---|---|---|
| 1 | 94 | 4 min | 1 |
| 2 | 94 | 30 sec | 10 |
| 3 | 65; -0.5 decrease per cycle | 30 sec | |
| 4 | 68 | 30 sec | |
| 5 | 94 | 30 sec | 28 |
| 6 | 60 | 30 sec | |
| 7 | 72 | 30 sec | |
| 8 | 72 | 7 min | 1 |
| 9 | 4 | Infinite Hold | -- |

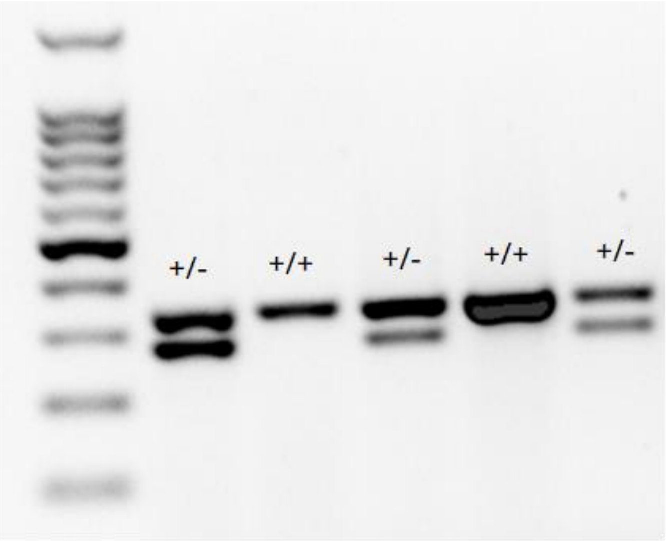


**Supplemental Figure 1. PCR genotyping to identify eNOS$^{+/-}$ fetuses following gel electrophoresis in a 1.7% agarose gel.**
Visualization was done using the Bio-Rad Gel Doc™ EZ Imager (Bio-Rad). eNOS$^{+/-}$ fetuses are indicated with +/- symbols showing two DNA products of ~300 bp and 337 bp. Wild-type fetuses show one DNA product of 337 bp and are indicated by symbols +/+.

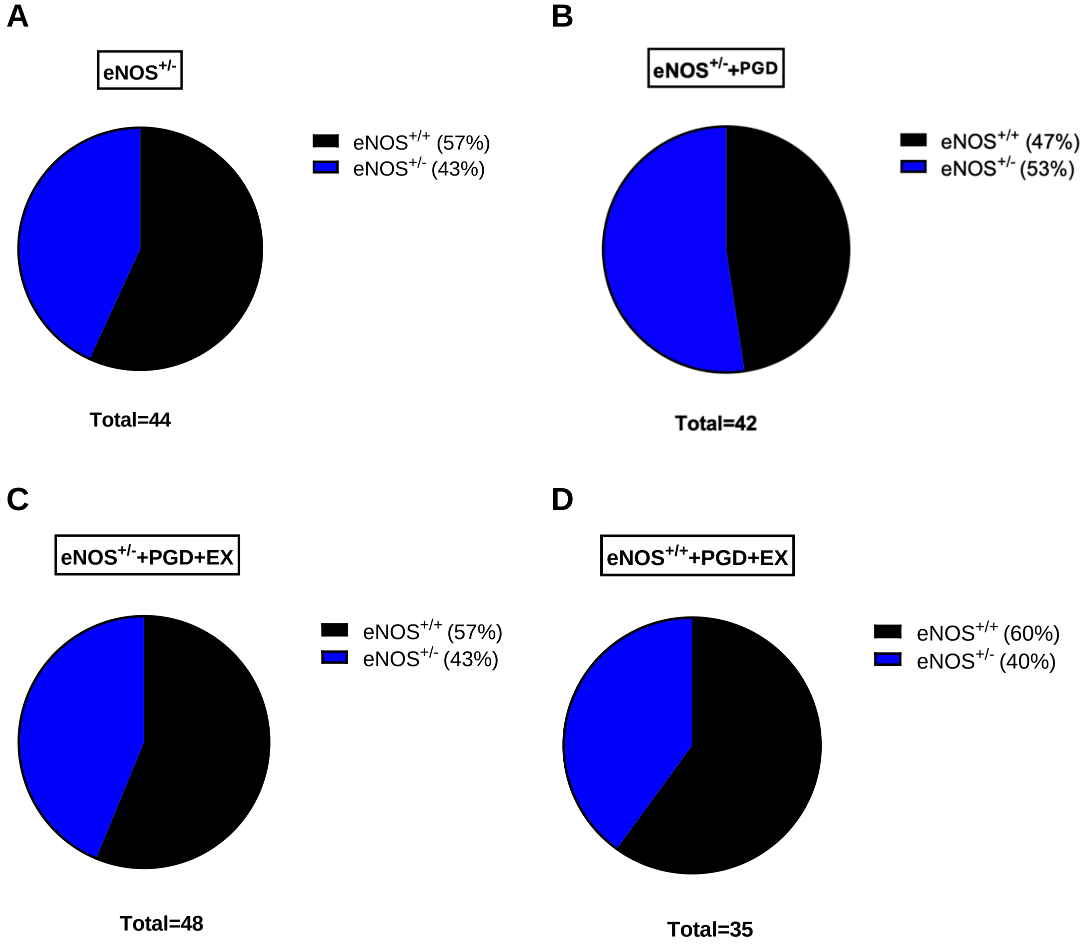


**Supplemental Figure 2. eNOS$^{+/+}$ and eNOS$^{+/-}$ litter proportions in each maternal condition determined via end point PCR.**

Litters from **(A)** eNOS$^{+/-}$ dams, **(B)** eNOS$^{+/-}$ dam with PGD, **(C)** eNOS$^{+/-}$ dams with PGD and maternal exercise (EX), **(D)** eNOS$^{+/+}$ dams with PGD and maternal exercise bred with healthy eNOS$^{+/-}$ males. Heterozygosity proportions all hovered around 50% which was expected. The proportion of eNOS$^{+/+}$ to eNOS$^{+/-}$ fetuses was found to be the same in each maternal condition (Chi square test $P$>0.05). eNOS: endothelial nitric oxide synthase, +/-: heterozygous, PGD: pregestational diabetes, EX: exercise.

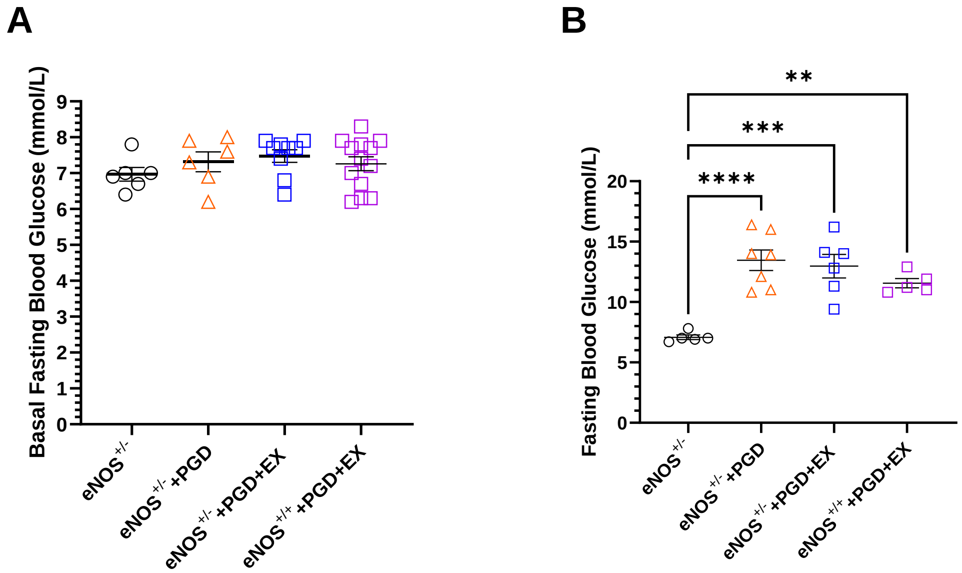


**Supplemental Figure 3. Basal 4 hour fasting blood glucose of dams before and after streptozotocin injection (PGD).**

**A)** No significant difference was found between groups before streptozotocin injection. **B)** Fasting blood glucose in dams was significantly elevated one week after STZ injection compared to control before pregnancy. Data are mean ± SEM of n=5–12 mice per group and were analyzed using one-way ANOVA followed by Tukey's test. ***P*<0.01, ****P*<0.001, *****P*<0.0001. eNOS: endothelial nitric oxide synthase, +/-: heterozygous, PGD: pregestational diabetes, EX: exercise.

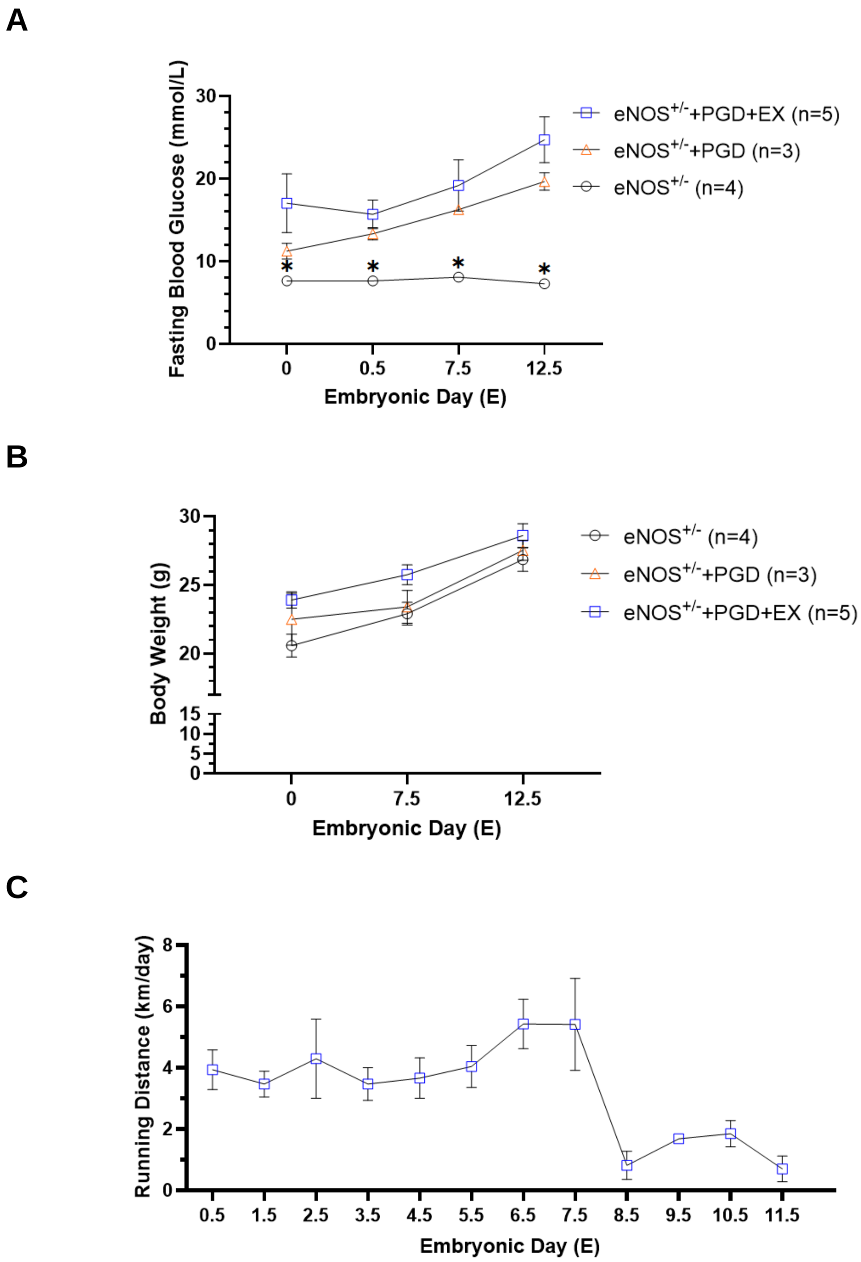


**Supplemental Figure 4. Maternal fasting blood glucose, body weight and running distance of eNOS$^{+/-}$ dams used for embryonic day 12.5 fetal collections.**

**A)** PGD conditions showed significantly elevated blood glucose levels compared to control over the entirety of gestation. Maternal exercise does not alter fasting blood glucose levels, as demonstrated by the similar fasting blood glucose observed in non-exercising PGD dams and exercising PGD dams. **B)** Maternal body weight was not impacted by PGD or voluntary exercise. **C)** Running distance by eNOS$^{+/-}$ PGD dams. Data represent mean ± SEM and were analyzed by one or two-way ANOVA followed by Tukey's test. *$P$<0.05 vs the corresponding data points of other 2 groups in A. eNOS: endothelial nitric oxide synthase, +/-: heterozygous, PGD: pregestational diabetes, EX: exercise.

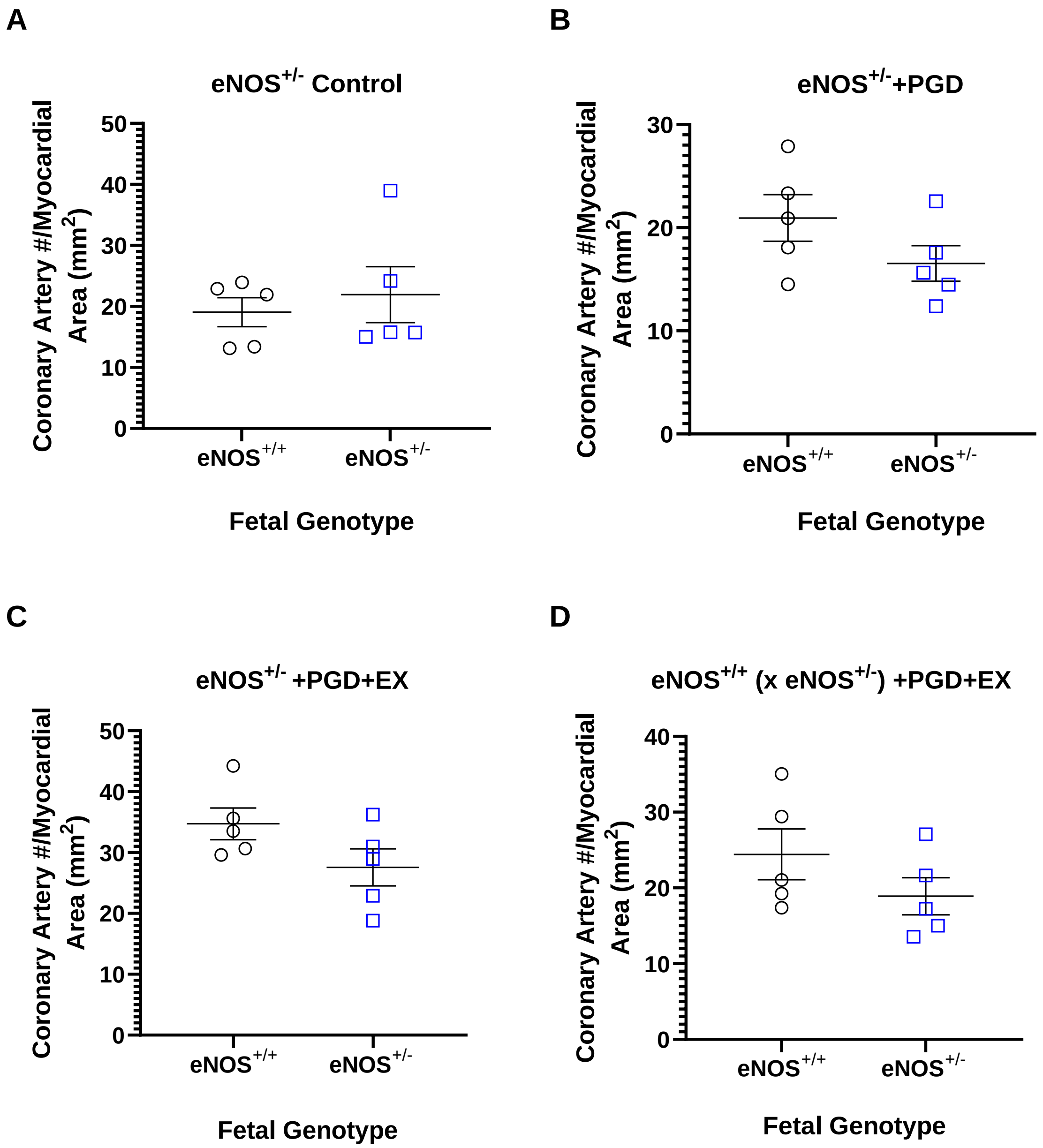


**Supplemental Figure 5. Coronary artery abundance relative to myocardial area in E18.5 eNOS$^{+/+}$ and eNOS$^{+/-}$ fetuses exposed to PGD with or without maternal exercise.**
**A)** eNOS$^{+/-}$ control dams. **B)** eNOS$^{+/-}$ dams with PGD. **C)** eNOS$^{+/-}$ dams with PGD and maternal exercise. **D)** eNOS$^{+/+}$ dams with PGD and maternal exercise bred with eNOS$^{+/-}$ males. The number of coronary arteries relative to myocardial area (mm²) was similar between eNOS$^{+/+}$ and eNOS$^{+/-}$ fetuses under each maternal condition. Data represents mean ± SEM of n=5 per group and were analyzed using unpaired Student's T-test, *P*> 0.05. eNOS: endothelial nitric oxide synthase, +/-: heterozygous, PGD: pregestational diabetes, EX: exercise.

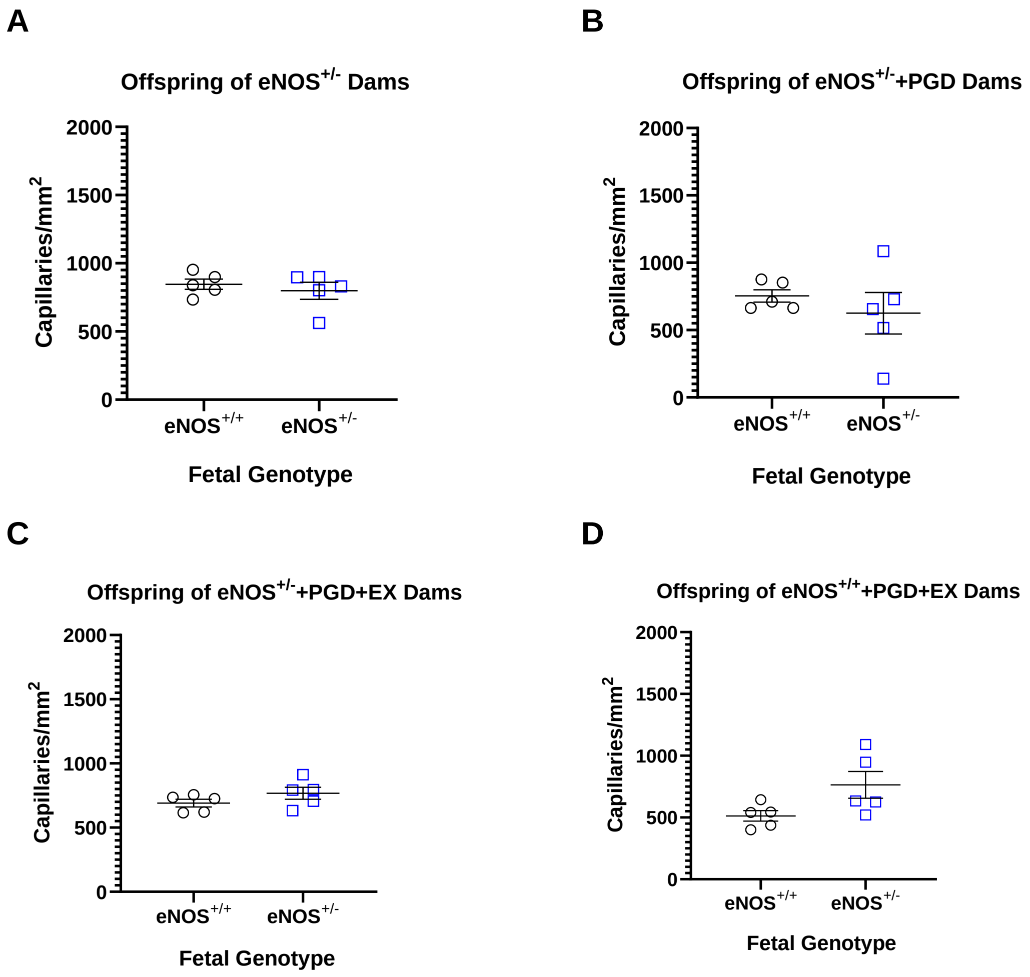


**Supplemental Figure 6. Capillary density relative to myocardial area in E18.5 eNOS$^{+/+}$ and eNOS$^{+/-}$ fetuses exposed to PGD with or without maternal exercise.**

**A)** eNOS$^{+/-}$ control dams. **B)** eNOS$^{+/-}$ dams with PGD. **C)** eNOS$^{+/-}$ dams with PGD and maternal exercise. **D)** eNOS$^{+/+}$ dams with PGD and maternal exercise bred with eNOS$^{+/-}$ males. Capillary density was similar between eNOS$^{+/+}$ and eNOS$^{+/-}$ fetuses exposed to PGD with or without maternal exercise. Data represent mean ± SEM of n=5 per group and were analyzed using unpaired Student's T-test, $P>0.05$. eNOS: endothelial nitric oxide synthase, +/-: heterozygous, PGD: pregestational diabetes, EX: exercise.

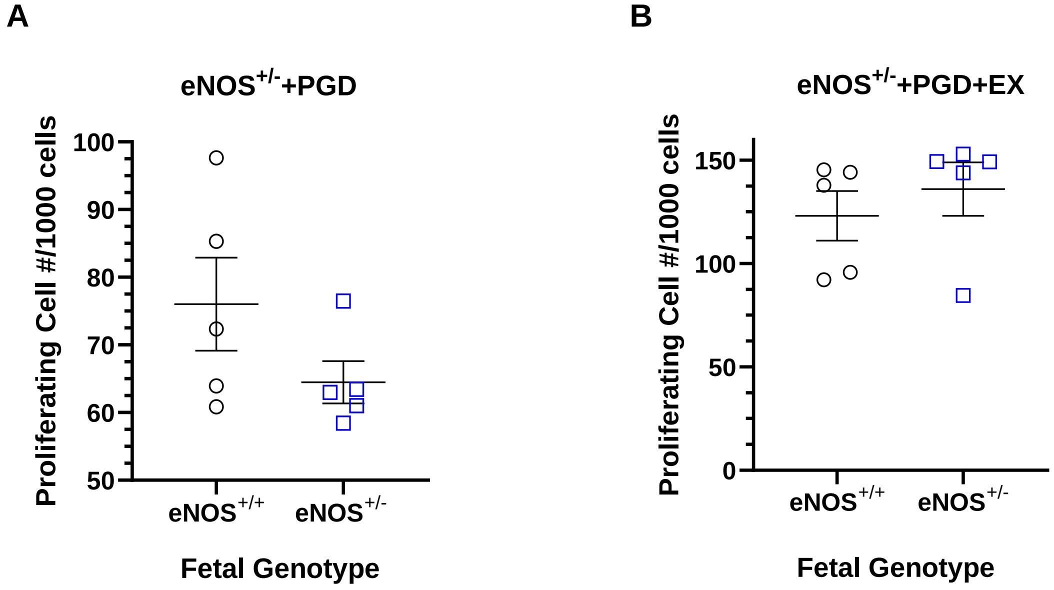


**Supplemental Figure 7. Number of proliferating cells in E12.5 eNOS$^{+/+}$ and eNOS$^{+/-}$ fetal hearts exposed to PGD with or without maternal exercise.**

**A)** E12.5 fetal hearts from eNOS$^{+/-}$ dams with PGD. **B)** E12.5 fetal hearts from eNOS$^{+/-}$ dams with PGD and maternal exercise. The proportion of proliferating cells was not significantly affected by pup genotype. Data represent mean ± SEM of n=5 per group and were analyzed by unpaired Student's T-test, $P>0.05$. eNOS: endothelial nitric oxide synthase, +/-: heterozygous, PGD: pregestational diabetes, EX: exercise

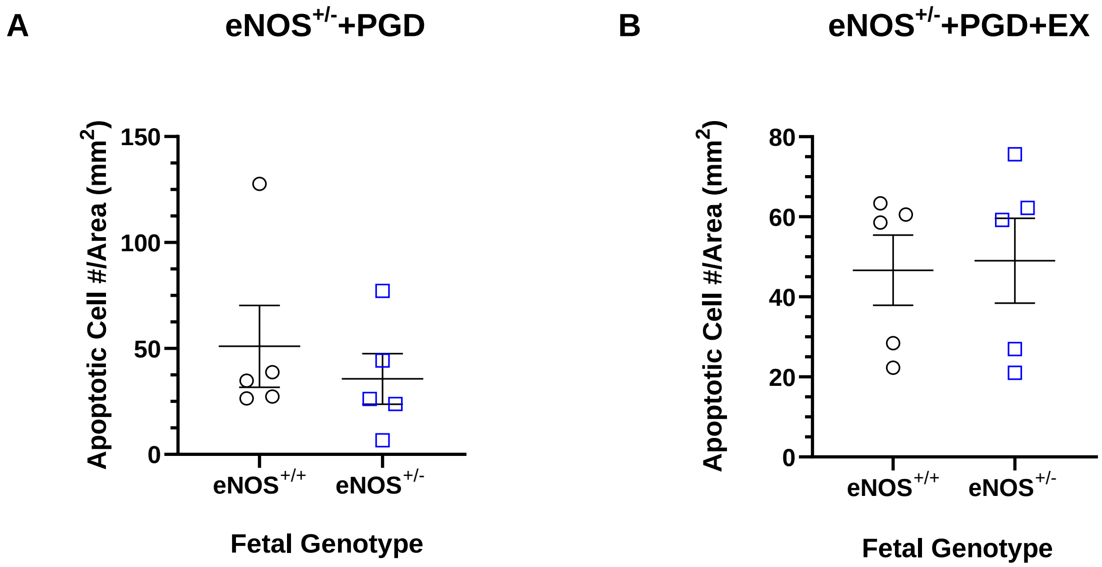


**Supplemental Figure 8. Number of cells undergoing apoptosis per area in E12.5 $eNOS^{+/+}$ and $eNOS^{+/-}$ mouse hearts exposed to PGD with or without maternal exercise.**

**A)** E12.5 fetal hearts from $eNOS^{+/-}$ dams with PGD. **B)** E12.5 fetal hearts from $eNOS^{+/-}$ dams with PGD and maternal exercise. The number of apoptotic cells per area ($mm^2$) was not significantly affected by pup genotype when exposed to PGD with or without maternal exercise. Data represent mean ± SEM of n=5 per group and were analyzed by unpaired Student's T-test, $P$>0.05. eNOS: endothelial nitric oxide synthase, +/-: heterozygous, PGD: pregestational diabetes, EX: exercise

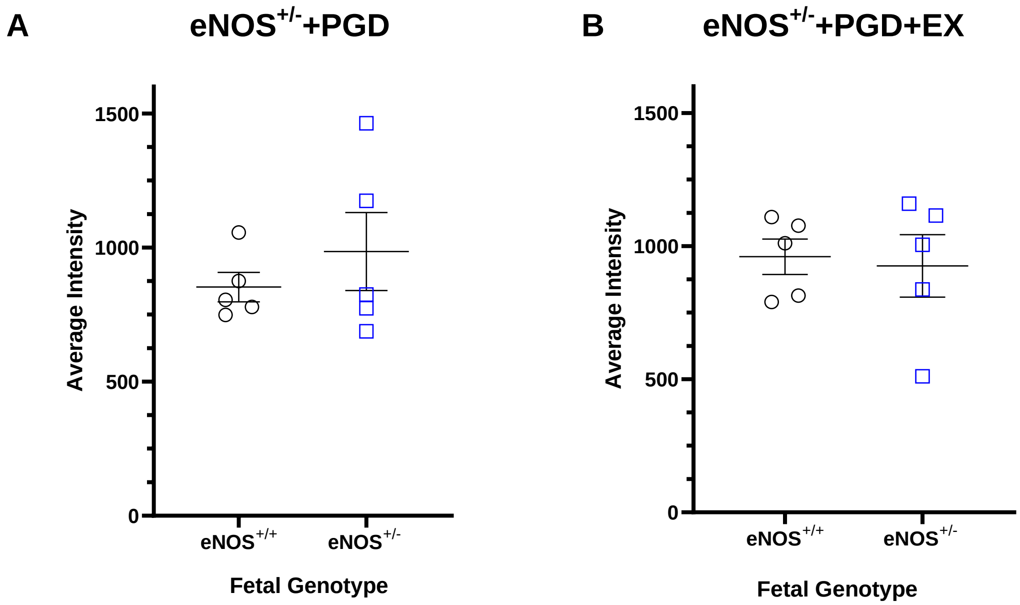


**Supplemental Figure 9. Superoxide levels, visualized using dihydroethidium (DHE), were similar between eNOS$^{+/+}$ and eNOS$^{+/-}$ fetuses exposed to PGD with or without maternal exercise.** DHE staining intensity in the hearts of fetuses exposed to **(A)** eNOS$^{+/-}$ dams with PGD and **(B)** eNOS$^{+/-}$ dams with PGD and maternal exercise. eNOS$^{+/+}$ and eNOS$^{+/-}$ fetuses did not influence superoxide levels exposed to PGD with or without maternal exercise. Data represent means ± SEM of n=5 per group and were analyzed by unpaired Student's T-test, $P$>0.05. eNOS: endothelial nitric oxide synthase, +/-: heterozygous, PGD: pregestational diabetes, EX: exercise

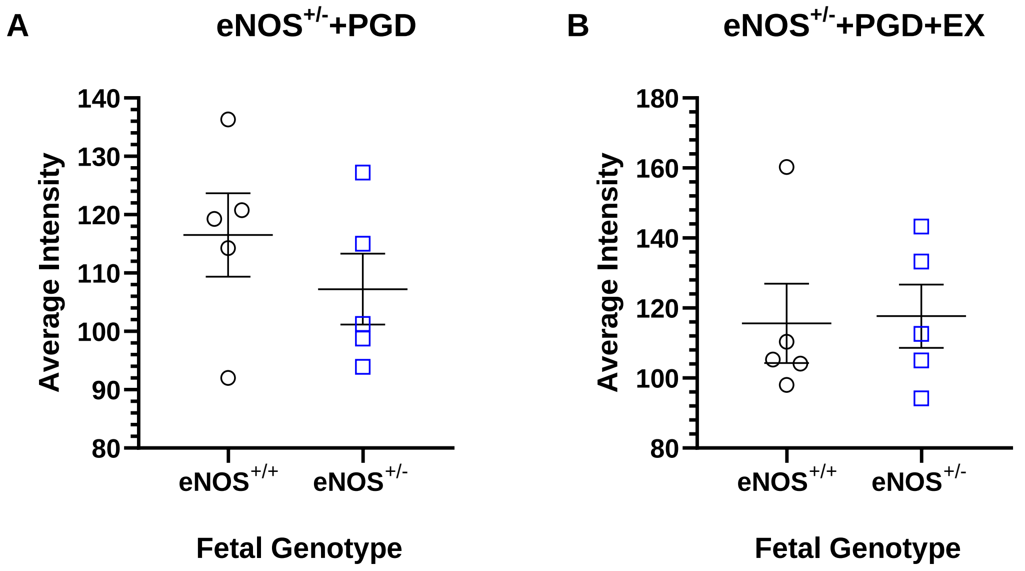


**Supplemental Figure 10. Lipid peroxidation levels visualized using 4-hydroxynonenal (4-HNE) levels were similar between eNOS$^{+/+}$ and eNOS$^{+/-}$ fetuses exposed to PGD with or without maternal exercise.**

4-HNE staining intensity in fetuses exposed to (A) eNOS$^{+/-}$ dams with PGD and (B) eNOS$^{+/-}$ dams with PGD and maternal exercise. eNOS$^{+/+}$ and eNOS$^{+/-}$ fetal genotypes did not influence superoxide levels exposed to PGD with or without maternal exercise. Data represent mean ± SEM of n=5 per group and were analyzed by unpaired Student's T-test, *P*>0.05. eNOS: endothelial nitric oxide synthase, +/-: heterozygous, PGD: pregestational diabetes, EX: exercise

**Supplemental Table 3. Incidence of PGD-induced CHDs according to severity.**

| **Maternal Condition** | **eNOS$^{+/-}$ (5 litters)** | | **eNOS$^{+/-}$ + PGD (7 litters)** | | **eNOS$^{+/-}$+PGD+EX (6 litters)** | | **eNOS$^{+/+}$ +PGD+EX (5 litters)** | |
|---|---|---|---|---|---|---|---|---|
| **Less Severe** | 5 | | 39 | | 27 | | 26 | |
| **More Severe** | 0 | | 1 | | 4 | | 4 | |
| **Fetal Genotype** | eNOS$^{+/+}$ (n=24) | eNOS$^{+/-}$ (n=20) | eNOS$^{+/+}$ (n=21) | eNOS$^{+/-}$ (n=21) | eNOS$^{+/+}$ (n=28) | eNOS$^{+/-}$ (n=22) | eNOS$^{+/+}$ (n=21) | eNOS$^{+/-}$ (n=14) |
| **Less Severe** | 1 | 4 | 18 | 21 | 17 | 10 | 14 | 12 |
| **More Severe** | 0 | 0 | 1 | 0 | 3 | 1 | 3 | 1 |

Fisher's exact test *P*>0.05. eNOS: endothelial nitric oxide synthase, +/-: heterozygous, PGD: pregestational diabetes, EX: exercise
